# scanMiR: a biochemically-based toolkit for versatile and efficient microRNA target prediction

**DOI:** 10.1101/2021.06.16.448293

**Authors:** Michael Soutschek, Fridolin Gross, Gerhard Schratt, Pierre-Luc Germain

## Abstract

microRNAs are important post-transcriptional regulators of gene expression, but the identification of functionally relevant targets is still challenging. Recent research has shown improved prediction of microRNA-mediated repression using a biochemical model combined with empirically-derived k-mer affinity predictions. Here, we translate this approach into a flexible and user-friendly bioconductor package, scanMiR, also available through a web interface. Using lightweight linear models, scanMiR efficiently scans for binding sites, estimates their affinity, and predicts aggregated transcript repression. Moreover, flexible 3’-supplementary alignment enables the prediction of unconventional interactions, such as bindings potentially leading to target-directed microRNA degradation (TDMD) or slicing. We showcase scanMiR through a systematic scan for such unconventional sites on neuronal transcripts, including lncRNAs and circRNAs.

## BACKGROUND

Since their initial discovery in *C.elegans*, microRNAs (miRNAs) have been shown to act on a plethora of biological pathways in diverse eukaryotic lineages (Bartel, 2018). They derive from short stem-loop precursor sequences, and after maturation associate with Argonaute (Ago) proteins to form the RNA-induced silencing complex (RISC) (Bartel, 2004). As part of this complex, miRNAs bind predominantly to the 3’ untranslated region (UTR) of mRNAs to induce post-transcriptional silencing (Jonas & Izaurralde, 2015). In animals, post-transcriptional silencing primarily comprises a combination of translational inhibition and mRNA degradation, with only a few miRNA target transcripts undergoing endonucleolytic cleavage (“slicing”) (Ameres & Zamore, 2013). Notable exceptions are target RNA-directed miRNA-degradation (TDMD) sites, which lead to a degradation of the bound miRNA (de la Mata et al., 2015; Fuchs Wightman et al., 2018; Han et al., 2020; Kleaveland et al., 2018; Shi et al., 2020). Binding specificity is determined in particular by the 5’ end of the miRNA, via the initial pairing of the so-called seed region that encompasses miRNA nucleotides 2-7 (Bartel, 2009), which can sometimes then propagate throughout the 3’part of the miRNA (Schirle et al., 2014; Sheu-Gruttadauria et al., 2019). The degree of repression that a miRNA imposes on its targets is determined by the time the RISC spends at individual sites, hence by the affinity of a miRNA for its mRNA targets (Denzler et al., 2016; McGeary et al., 2019). Strikingly, due to the influence of Ago on the functional properties of this interaction, the binding kinetics do not resemble conventional RNA:RNA hybridization (Salomon et al., 2015), making it difficult to predict binding efficiency through traditional computational methods.

To date, several bioinformatic approaches have been proposed to identify functional miRNA targets and to approximate their repression efficiency (e.g. TargetScan 7 (Agarwal et al., 2015), DIANA-microT-CDS (Reczko et al., 2012), MIRZA-G (Gumienny & Zavolan, 2015), and MirTarget (Liu & Wang, 2019); see Kern et al. (2020) for a comprehensive review). Most of these methods follow correlative approaches in that they try to infer a repression score from a number of features shown to correlate with repression efficiency of miRNAs, for example structural accessibility, predicted pairing stability, local AU-content, site-type, and 3’UTR length (Agarwal et al., 2015; Grimson et al., 2007; Gumienny & Zavolan, 2015; Krek et al., 2005; Nielsen et al., 2007). Some of these algorithms include the probability of sequence conservation since it has been shown that some of the parameters mentioned before are enriched in the proximity of evolutionarily conserved sites (Grimson et al., 2007; Nielsen et al., 2007). One of the publicly-available prediction tools most accurate in explaining observed repression in cellular contexts (TargetScan7) has been reported to perform as well as recent CLIP experiments (Agarwal et al., 2015). Nevertheless, TargetScan7 still explains only a small fraction (~14%) of the variance in logFC in cells perturbed with miRNA mimics (Agarwal et al., 2015).

Recently, several high-throughput studies have attempted to shed light on the biochemical determinants of miRNA-mRNA targeting (Becker et al., 2019; McGeary et al., 2019, 2021; Salomon et al., 2015). In one of these studies, the investigators used purified Ago-miRNA complexes in combination with RNA bind-n-seq (RBNS) to determine the affinities of six miRNAs (miR-1, let-7a, miR-7, miR-124, miR-155 and lsy-6, selected based on their conserved sequence and existing functional data), to any possible 12 nt long (12-mer) sequence (McGeary et al., 2019). They showed that the affinities and the corresponding dissociation constants (log(K_D_)-values) of different canonical site-types correlate very well with repression observed in cellular contexts, substantially outperforming a retrained version of TargetScan7. In an attempt to extrapolate these findings to other miRNAs, the authors trained a convolutional neural network (CNN) using the empirically obtained data. They demonstrated that repression scores based on affinity values obtained via the CNN outperform TargetScan7 by ~50% across 12 different miRNAs in another cell-type (McGeary et al., 2019).

While the model and experimental data provided by McGeary et al. (2019) represent major advances in target prediction, they are not available to a broader class of users. Indeed, while occupancy scores based on the biochemical model have been added to TargetScan8, such widely used platforms fail to offer the needed flexibility: TargetScan for instance includes only a handful of species, covers only a subset of the transcriptome (single curated UTR per protein-coding gene), does not provide information on non-canonical or open reading frame (ORF) binding sites and is very tedious to apply to custom sequences. Finally, despite the prevalence of R and Bioconductor in bioinformatics, there is a relative paucity of R-based miRNA target prediction tools. We therefore sought to address these needs by integrating and extending the biochemical model of McGeary et al. (2019) within the Bioconductor framework to offer a generally applicable, flexible, and user-friendly package and accompanying web app, *scanMiR* (Fig. 1). *scanMiR* efficiently scans for canonical and noncanonical miRNA binding sites on any custom sequence (incl. circular RNAs), estimates dissociation constants and predicts aggregated transcript repression. We further include a flexible 3’-supplementary alignment, allowing the systematic prediction of putative TDMD sites as well as sites leading to RNA slicing. Moreover, *scanMiR* provides several novel visualization functions in order to facilitate miRNA-target candidate selection. *scanMiR*-predicted mRNA repression correlates significantly better with measured mRNA fold-changes than TargetScan scores. Finally, transcript-level predictions for human, mouse and rat transcriptomes, as well as the visualization and scanning functions of *scanMiR*, are available to a broader public through a user-friendly webtool.

**Figure 1:**
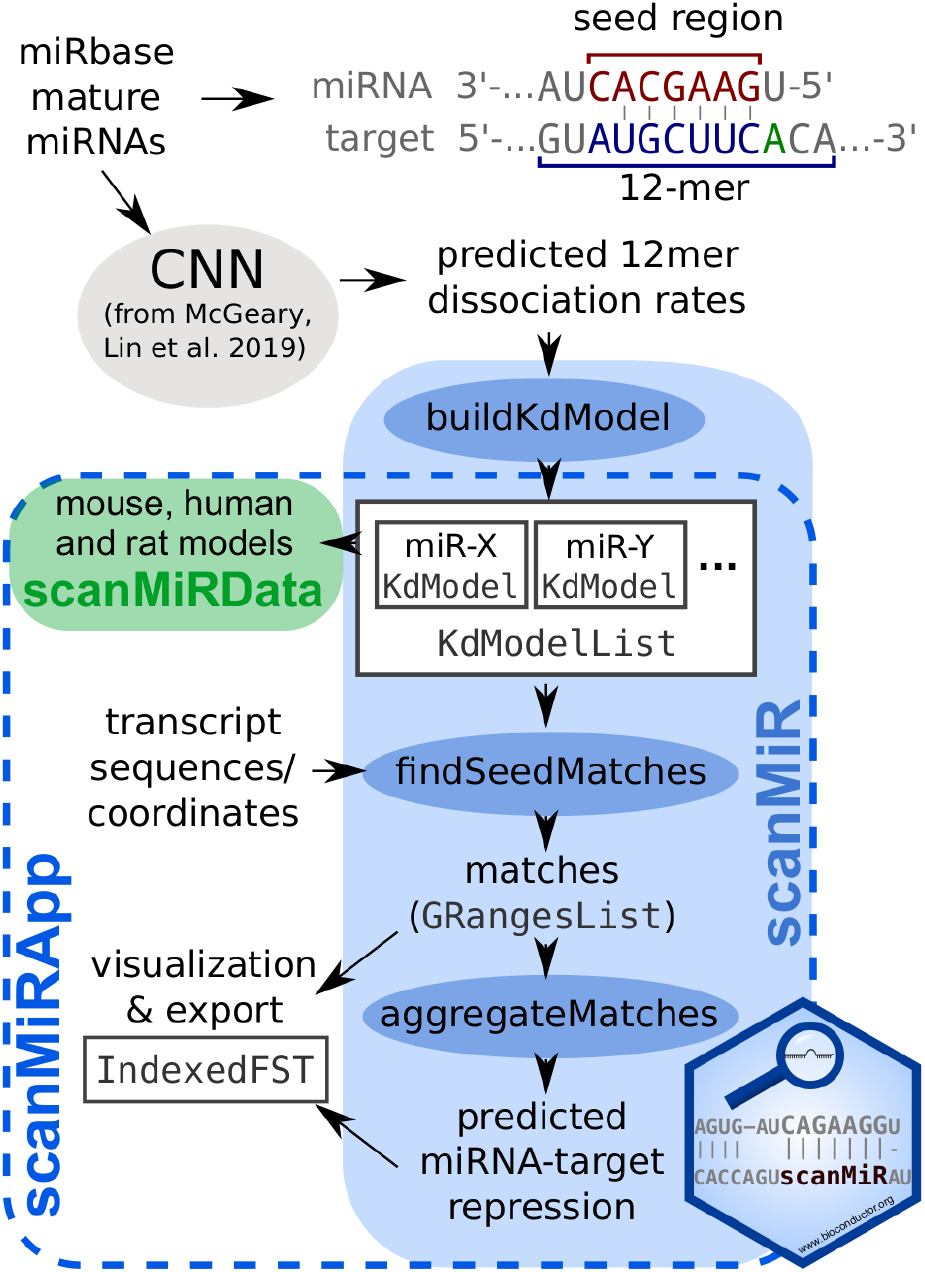
Overview of the *scanMiR* suite. The scanMiR package offers classes and functions to generate and work with miRNA 12-mer affinity models (i.e. 8nt opposite the extended seed region plus flanking dinucleotides). It enables efficient and flexible site scanning, as well as different methods to visualize alignments. The *scanMiRData* package contains predicted affinity models for all human, mouse and rat miRbase miRNAs. Finally, the *scanMiRApp* package provides an easily deployable web interface to most *scanMiR* functionalities, as well as convenient wrappers to facilitate data handling and integration with annotation packages.

## RESULTS

### Efficient encoding of miRNA dissociation constants

McGeary, Lin and colleagues (2019) developed a convolutional neural network (CNN) predicting the dissociation constants (K_D_) of given miRNAs on partially-matching 12-nucleotides sequences (that is, the 8nt opposite the extended miRNA seed sequence as well as two flanking di-nucleotides on each site, see the scheme in Fig. 1 and Methods for further infos). Given that there are about half a million possible 12-mers and corresponding K_D_s for each miRNA, it is impractical to store and use the information across all possible miRNAs. We therefore developed an efficient compression and encoding of the K_D_ predictions (see Methods), enabling its rapid application for scanning candidate target transcripts. As Figure 2A-B shows, the correlation of the reconstituted K_D_-values to the original ones is high, and above the accuracy of the CNN predictions (McGeary et al., 2019). With this approach, we could for instance store the affinity models for ~2000 miRNAs in approximately 10 MB (instead of over 8 GB for the storage of 12mer-K_D_ pairs) and then apply them very efficiently for the identification and characterization of putative sites (Fig. 3D).

**Figure 2:**
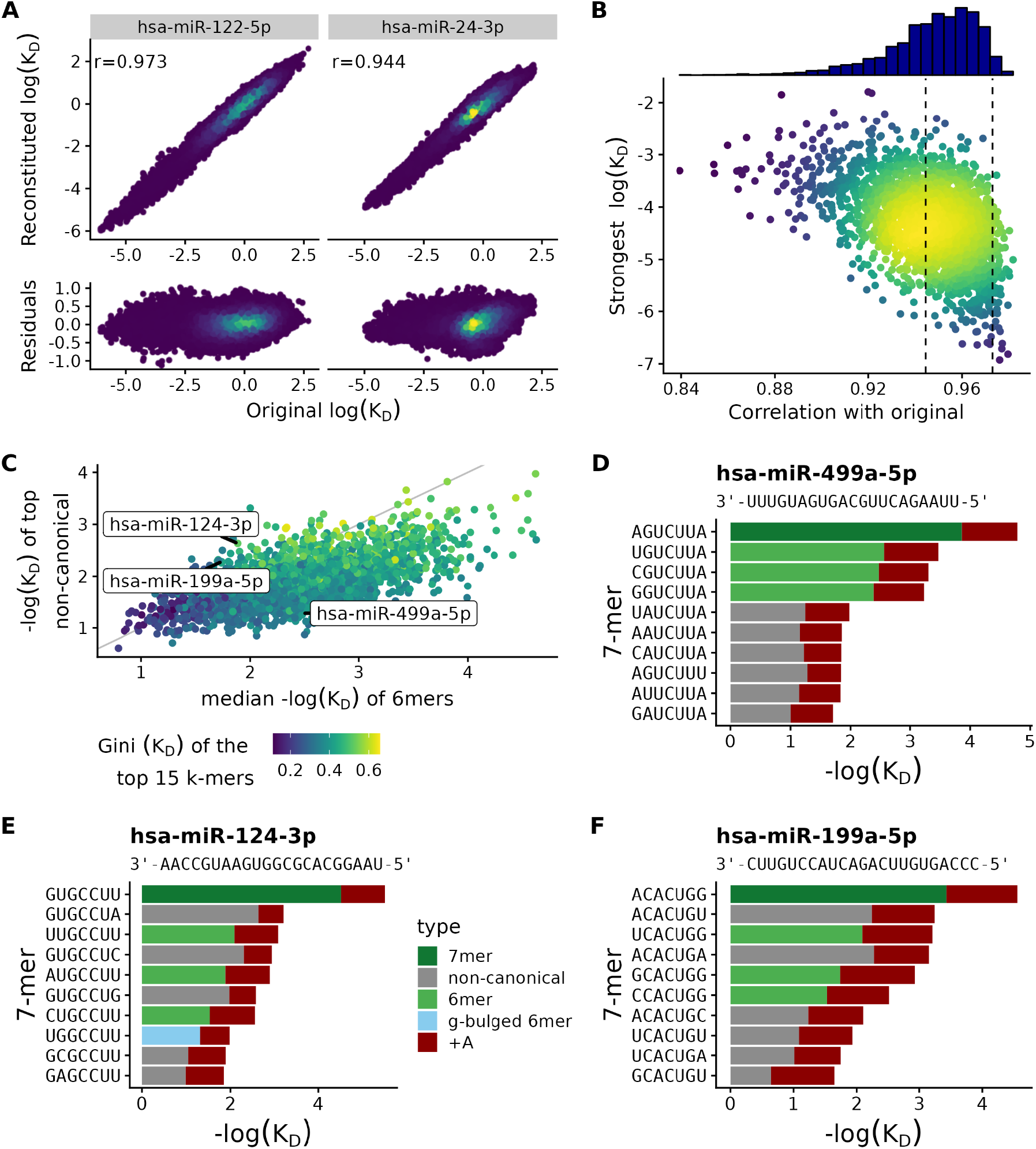
Efficient encoding of miRNA dissociation rates. **A:** Correlation between log(K_D_) values before and after compression for two example miRNAs (a known high-specificity miRNA, miR-122, as well as the miRNA closest to the mean correlation (see panel B). The residuals (reconstituted-original) are shown below. **B:** Distribution of correlations across all human mirBase putative miRNAs. The dashed lines represent the two miRNAs selected in panel A. The miRNAs with a poor correlation tend to be those with weak dissociation rates, hence spanning a smaller dynamic range, indicating that this is less an effect of the compression than of the poor predicted specificity of the miRNA. **C:** Relationship between the dissociation constants of the 6-mers sites and that of the top non-canonical sites, highlighting miRNAs with radically different binding behaviors. miRNAs are colored according to the Gini coefficient across the K_D_ of their top 15 k-mers. **D-F:** Example affinity plots showing the top ten miRNA seed complements ranked by the average dissociation constant for the miRNA of interest. MiR-499a-5p displays a very specific canonical binding (**D**), whereas miR-124-3p and miR-199a-5p include highly ranked non-canonical seeds (**E-F**).

**Figure 3:**
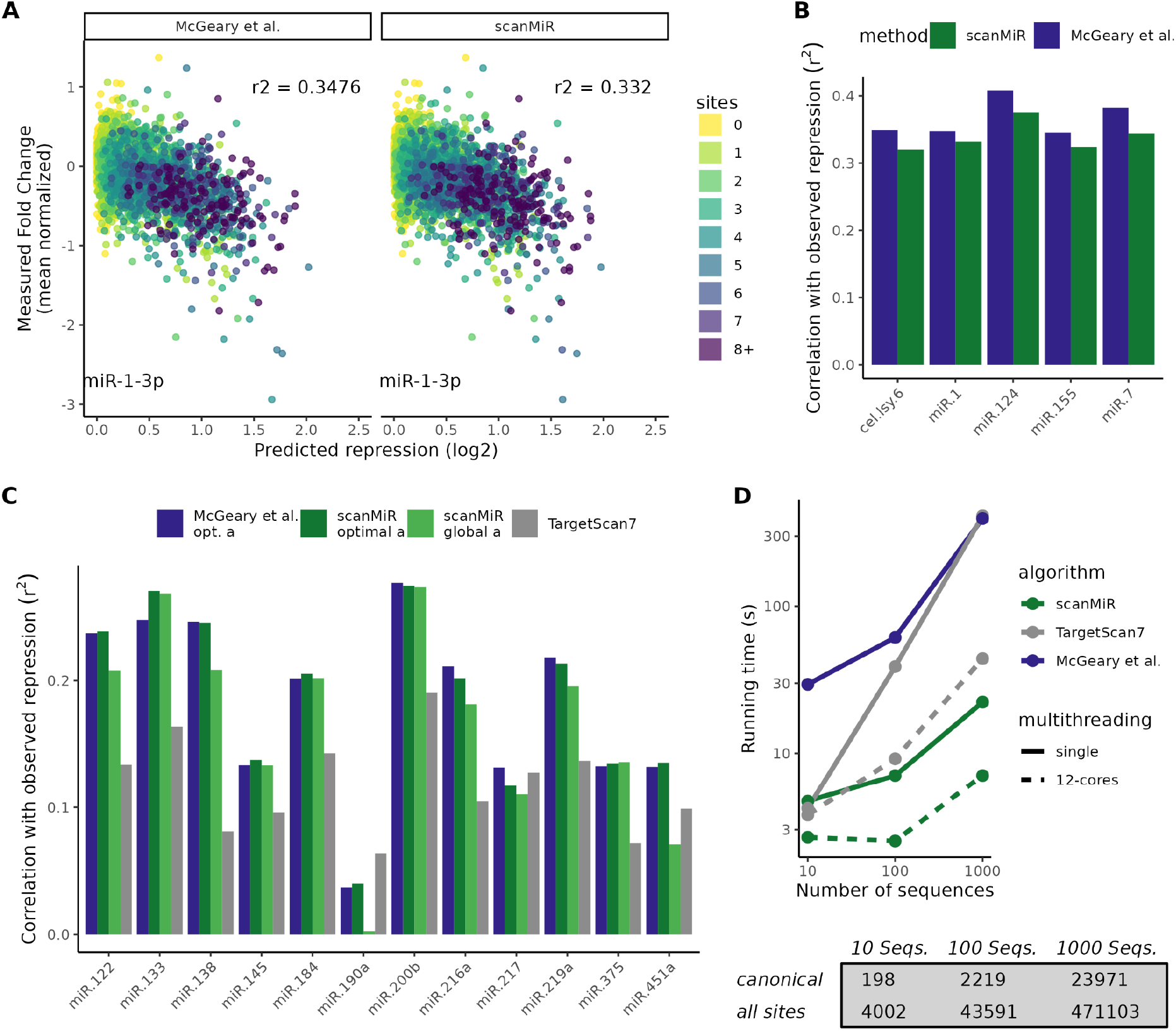
Efficient prediction of repression. **A:** Correlation between predicted and observed repression in an example miRNA (hsa-miR-1-3p) transfection experiment from McGeary et al. (2019), showing comparable correlations between the original algorithm and log(K_D_) values and our compressed version. Each point corresponds to a transcript, and is colored according to the number of canonical sites in its UTR. **B:** Correlation (Pearson *r^2^*) between predicted and observed repression across the 5 miRNAs from McGeary et al. (2019) with experimental affinity measurements. **C:** Correlation (Pearson *r^2^*) between predicted and observed repression across 12 further miRNAs in HEK cells (McGeary et al., 2019). The authors had chosen those particular 12 miRNAs because of their very low endogenous expression in HEK cells. Correlations obtained with a globally optimized *a*-value are shown in light green, correlations based on TargetScan7 predictions in grey. **D:** Running times of the different methods scanning for binding sites of 12 miRNAs across growing sets of sequences (the sites indicated are from scanMiR before filtering, including all canonical and non-canonical sites).

The *KdModel* and *KdModelList* classes defined in *scanMiR* provide an interface for handling these objects. We generated *KdModel* objects for all human, mouse and rat putative miRNAs from miRbase (Griffiths-Jones et al., 2006), which are made available in the accompanying *scanMiRData* package.

In an attempt to draw further biological conclusions from seed properties stored in KdModels, we set out to separate human miRNAs based on patterns of binding specificity (Fig. 2C). Specifically, we examined the relationship between the dissociation constants of the top non-canonical seed and the median of canonical 6mers (generally accounting for 7mer-A1 and 6mer sites). miRNAs did not cluster into distinct classes but roughly formed a multivariate normal distribution. Nevertheless, we could recognize different paradigmatic examples, such as the highly-specific binding of miR-499a-5p (Fig. 2D) and the known strong non-canonical binding of miR-124-3p (Chi et al., 2012; Wang et al., 2021) (Fig. 2E). Based on its proximity to miR-124-3p in the distribution, we likewise propose the occurrence of previously uncharacterized high-affinity non-canonical binding sites for miR-199a-5p (Fig. 2F).

An important experimental aspect of miRNA research is the mutation of predicted binding sites to investigate their putative function. Although it is known that nucleotide substitutions opposite miRNA positions 2-5 within the seed region most efficiently affect binding (Becker et al., 2019), the importance of single nucleotides within the seed region of different miRNAs has not yet been elucidated. By generating positional weight matrices of miRNAs based on the K_D_-predictions of 12mer sequences, *scanMiR* allows the user to directly plot the nucleotide information content of individual miRNAs. Such affinity based nucleotide information content plots suggest for example a uniform importance of nt 2-5 in the seed of miR-499a-5p, whereas for miR-129-5p, nt 3-4 seem to be most relevant for a stable interaction with its targets (Suppl. Fig. 1A). Researchers can easily resort to this information to get a first impression on which nucleotides in the seeds of miRNAs are most required for a stable interaction with their targets, and hence should be primarily mutated to prevent miRNA binding.

### Biochemically-based prediction of miRNA target repression

To assess the accuracy of compressed *KdModels* for miRNA repression prediction, we scanned HeLa mRNAs for putative binding sites of five miRNAs that McGeary and colleagues (2019) had initially used for their RBNS experiments and determined site affinity based on the compressed models (see Methods). Aggregating these values using the biochemical model and correlating them to mRNA fold-changes observed in Hela cells following miRNA-mimic transfections revealed similar results as with uncompressed dissociation constants (Fig. 3A-B, Suppl. Fig. 2C). This holds likewise true for 12 further transfection experiments that were conducted in another cell type (HEK293) with miRNAs that were not used in the direct RBNS-measurements (Fig. 3C). Importantly, for most of these miRNAs, the correlation values are considerably higher than those obtained with TargetScan7 repression scores (Fig. 3C). This also holds true in comparison to the occupancy scores recently made available in TargetScan8, which are likewise based on the biochemical model of McGeary et al. (2019), but are only provided for a subset of miRNAs on a set of pre-curated transcripts (Suppl. Fig. 2).

One of the constraints of the biochemical model is the necessity to fit the relative concentration of unbound AGO-miRNA complexes (*a*-value) for each miRNA and experiment individually (see Methods, (McGeary et al., 2019)). Bolstered by the observation that even a 100-fold deviation of the optimal *a*-value still leads to a repression prediction outperforming TargetScan7 (Suppl. Fig. 3B) (McGeary et al., 2019), we went on to generate a globally optimized *a*-value using the 12 HEK-cell transfection datasets. Repression predictions using this global value are slightly lower than those with *a* optimized separately for each dataset, but still outperform TargetScan substantially (Fig. 3C, Suppl. Fig. 2). To further assess the general validity of our approach, we tested predicted miRNA repression on published mouse miRNA knockout datasets. *scanMiR* similarly outperforms TargetScan in this experimental setup, although the overall signal-to-noise ratio and resulting explanatory power were lower (Fig. Suppl. 4). In conclusion, *scanMiR* repression predictions exceed the performance of the latest TargetScan7&8 scores across several experimental settings, and the repression predictions are roughly generalizable even without fitting the biochemical parameters to individual experimental settings.

A further key finding of McGeary et al. (2019) was the identification of high-affinity non-canonical miRNA binding sites. By harnessing the whole set of available 12mer sequences and corresponding K_D_-values to identify and rank miRNA binding sites, *scanMiR* readily detects these presumably efficient non-canonical sites. However, including also the vast number of non-canonical sites with low predicted target affinities significantly exacerbates the scanning and storage of miRNA binding sites. We therefore studied the relation between K_D_-values of different binding sites and their explanatory power. For 23 out of 24 miRNA-transfection experiments in HEK- and HeLA-cells with an explanatory power larger than 10%, a lenient log(K_D_)-cutoff value of −0.3 retained 97.5% of this power while at the same time allowing a reduction in the number of stored sites by over 40%, and a more stringent −0.5 cutoff also incurred very little additional loss in power (Suppl. Fig. 5A-B).

As an additional *scanMiR* feature, we implemented the possibility to flag g-bulged as well as wobbled (G:U) miRNA binding sites in the scanning results as well as in affinity plots (Fig. 2E, Suppl. Fig 1B-C). G-bulged non-canonical binding sites comprise a bulged out “G” opposing Nt6 of the miRNA to allow further seed pairing, and the efficient binding of, for example, miR-124-3p to such sites has been shown by high-throughput techniques and the functional repression of candidate targets validated with luciferase experiments (Chi et al., 2012; Wang et al., 2021).

Moreover, *scanMiR* enables the user to specify a minimum distance between binding sites. It has been shown previously that binding sites located closer than 7nt to each other exert a repelling effect onto each other (Grimson et al., 2007; Sætrom et al., 2007). By default, *scanMiR* therefore removes binding sites within the specified minimal distance of a higher-affinity site (Suppl. Fig. 6, see Methods).

Of note, despite evaluating a considerably larger number of sites (i.e. including all non-canonical sites) than TargetScan, *scanMiR* nevertheless runs significantly faster. It likewise runs faster than the scripts from McGeary et al. (2019) (Fig. 3D), which were even slower when including the default transcript folding (Suppl. Fig. 5D).

### Flexible 3’ supplementary alignment flags special miRNA binding sites

Upon finding a partial seed match, *scanMiR* performs a local alignment of the 3’ region of the miRNA onto the upstream region of the transcript, thereby enabling a flexible and customizable miRNA-mRNA bulge size (Suppl. Fig. 7A). Including a 3’-supplementary pairing score based on this alignment had minimal impact on *scanMiR* repression predictions (Suppl. Fig. 7B-C); though more importantly, it enables the prediction of unconventional binding sites determined by this supplementary pairing. It has been shown previously that some miRNA binding sites with near full-length complementarity can lead to endonucleolytic cleavage (“slicing”) of the target sequence (Carthew & Sontheimer, 2009). Experimentally validated examples comprise the slicing of HOXB8 upon binding of miR-196a-5p (Yekta et al., 2004) or the slicing of the CDR1as by miR-671-5p (Hansen et al., 2011; Kleaveland et al., 2018). *scanMiR* correctly identifies and represents these sites (Fig. 4A-C).

**Figure 4:**
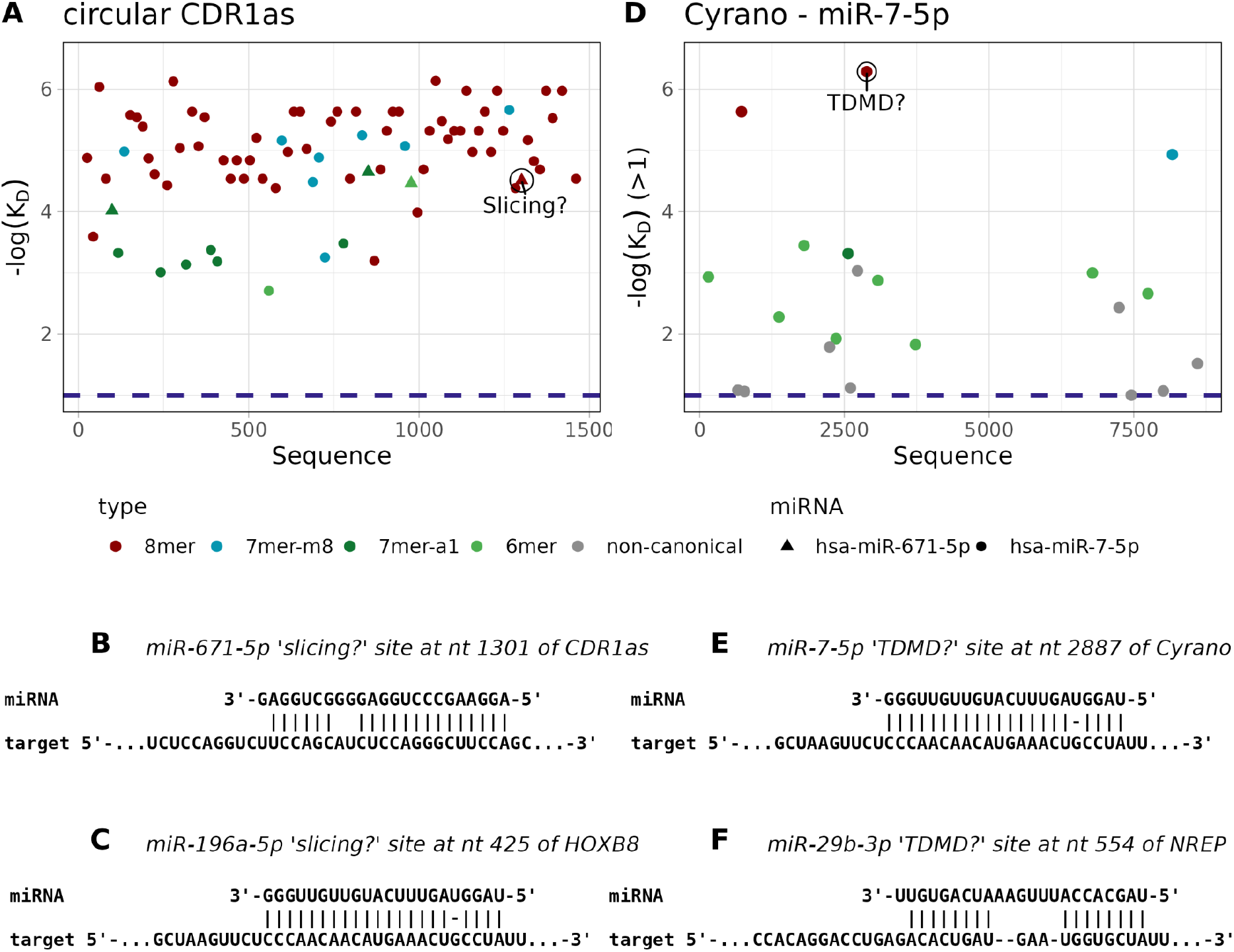
3’ supplementary alignment enables the prediction of special miRNA sites. **A:** MiR-7-5p and miR-671-5p binding sites on the CDR1as circular RNA, revealing especially a very large concentration of miR-7-5p sites. Binding of miR-671-5p to the circular RNA CDR1as leads to endonucleolytic cleavage of this circRNA. The binding site is flagged by scanMiR as “Slicing?” **B:** The full alignment of the validated miR-671-5p slicing site on CDR1as. **C:** Binding of miR-196a-5p to HOXB8 likewise leads to slicing of the target mRNA. scanMiR also flags this site as “Slicing?”, despite the low affinity of the seed of miR-196a-5p to HOXB8 (log(K_D_) = −1.27), which can be explained by the G:U wobble binding at position five. Shown is the full miRNA-mRNA alignment at this site. **D-E:** Validated miR-7-5p TDMD site on the Cyrano RNA and its alignment up to the last 3’ nucleotides of the miRNA. **F:** Full alignment of the likewise experimentally validated miR-29b-3p TDMD site on the NREP mRNA. All examples use human transcript and miRNA annotations.

Moreover, sufficient target complementary at the 3’ end of the miRNA in combination with specific mRNA-miRNA bulges can lead to target-directed miRNA degradation (TDMD) (Ameres et al., 2010; de la Mata et al., 2015). Specifically, the Ago-miRNA complex undergoes a structural change at TDMD sites, which enables a ubiquitin ligase (ZSWIM8) to bind and poly-ubiquitinate the complex, thereby ultimately leading to the degradation of the miRNA (Han et al., 2020; Sheu-Gruttadauria et al., 2019; Shi et al., 2020). By taking these determinants into account (see Methods), *scanMiR* flags possible TDMD sites, as shown for the experimentally validated examples of miR-7-5p binding to the lncRNA Cyrano (Fig. 4D-E) (Kleaveland et al., 2018) and the miR-29b-3p site on the mRNA of NREP (Fig. 4F) (Bitetti et al., 2018).

Based on these results and the recent finding that TDMD-induced miRNA turnover might be more prevalent in different cell types than previously anticipated (Han et al., 2020; Shi et al., 2020), we decided to analyse the occurrence of potential TDMD sites more systematically. We focussed on miRNAs that were significantly upregulated in induced mouse neurons upon ZSWIM8 knockout, suggesting that they undergo TDMD in this cell type (Shi et al., 2020). To get an idea of which binding sites might trigger the degradation of these miRNAs, we obtained the expression values of transcripts with predicted TDMD sites from induced mouse neurons at the same day of differentiation (Whipple et al., 2020). This analysis revealed again the well expressed known TDMD sites for miR-7a-5p on Cyrano and miR-29b-3p on Nrep, but also several other interesting candidates, for example, another potential TDMD site for miR-7a-5p on the highly expressed Dpysl3, as well as potential TDMD sites for miR-495-3p on two isoforms of the heat-shock protein Hsp90aa1 (Fig. 5A). In addition, we specifically screened sequences obtained from a recent full-length profiling of circRNA isoforms expressed in the mouse brain for the presence of TDMD sites of the aforementioned miRNAs (Zhang et al., 2021). This analysis suggests potential TDMD sites, for example, on the abundant circRNAs Mef2a and Akt3 for miR-297c-5p or miR-376b-3p, respectively (Fig. 5B). In conclusion, our analyses suggest several candidate mRNAs, lncRNAs and circRNAs that might be involved in the regulation of miRNA abundance (Supplementary Tables 1-2).

**Figure 5:**
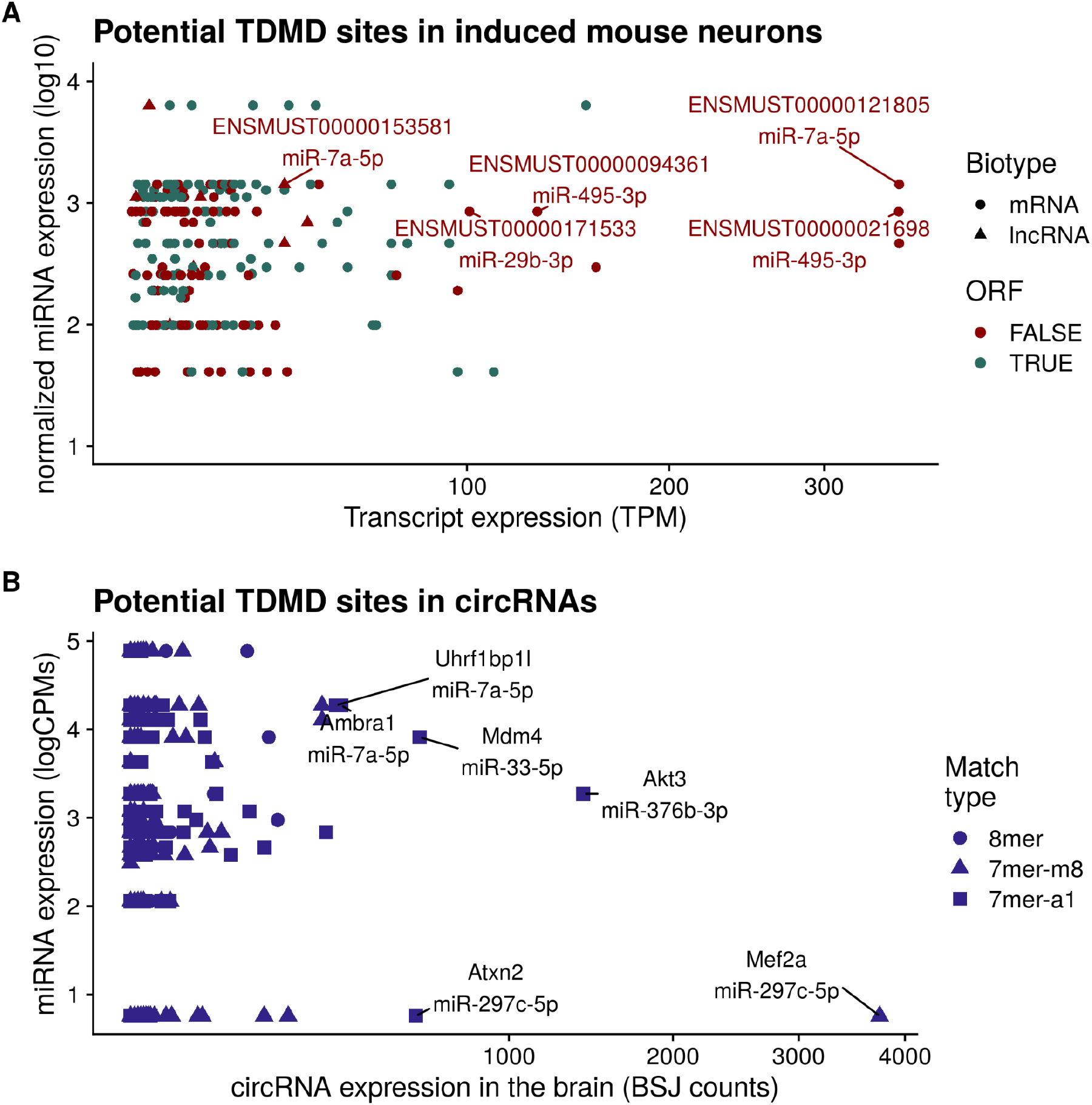
Prediction of potential TDMD sites in neurons. Potential TDMD sites of miRNAs significantly upregulated in induced mouse neurons upon ZSWIM8 knockout (Shi et al., 2020). **A:** scanMiR predicted TDMD sites on expressed linear transcripts as obtained from an RNA sequencing by Whipple et al., (2020). The normalized miRNA expression in wt-neurons from Shi et al. (2020) is plotted on the y-axis. **B:** scanMiR predicted TDMD sites on circular RNAs expressed in the mouse brain (Zhang et al., 2021). miRNA expression values (plotted on the y-axis) are taken from Chiang et al. (Chiang et al., 2010).

### Prediction of circRNA sponges/transporters in the brain

Due to their stability, it has been hypothesized that circular RNAs (circRNAs) with multiple miRNA binding sites could act as sponges or transporters (Kristensen et al., 2019). The prevalence of such mechanisms however remains unknown. To predict circRNAs that might exert these roles in the mammalian brain, we identified circRNAs that are significantly enriched for canonical binding sites of specific miRNAs using a binomial model conditioned on the total number of binding sites for the given miRNA on each circRNA and the number of all other miRNA binding sites on the given circRNA as background (see methods). Reasoning once more that lowly-expressed circRNAs are unlikely to have significant biological impact, we plotted this enrichment against the abundance of the circRNAs (Suppl. Fig. 8A). MiR-7a-5p binding sites on the (by far) most abundant circRNA Cdr1as were most significantly enriched, in line with the previously characterized exceptional effectiveness of Cdr1as in binding this miRNA (Kristensen et al., 2019). Nevertheless, our data shows that several additional highly-expressed circRNAs contain significantly more putative miRNA binding sites than expected by chance, suggesting a potential functional role (see also Supplementary Table 3).

Finally, it has been proposed that miRNA-mediated cleavage might enhance circRNA turnover rates and result in the release of bound miRNAs as well as RNA-binding proteins (Kristensen et al., 2019). To investigate this hypothesis, we conducted an additional screen for slicing sites on circRNAs. This analysis retrieved only a few candidates (Suppl. Fig. 8B, Supplementary Table 4), among them the known Cdr1as, suggesting that miRNA-dependent slicing plays a minor role in the context of circRNA biology.

### A web interface to *scanMiR*

To make *scanMiR* accessible to a broader public, we developed a web interface to its main functionalities, available at https://ethz-ins.org/scanMiR/. The interface allows the visualization of mouse, human and rat *KdModels*, the interrogation of top targets for all miRbase miRNAs, the easy and flexible scanning of any transcript sequence for miRNA binding sites, and the visualization of binding sites including their supplementary 3’ alignment (Suppl. Fig. 9). The application can additionally be easily deployed by anybody through the *scanMiRApp* package, for instance enabling groups working on other species to compile their custom annotation, launch the app locally, or set up their own portals. *scanMiRApp* includes a number of wrappers (notably connecting with AnnotationHub to easily include updated transcript annotations), as well as features to enable rapid out-of-memory random access to large scans (see Methods).

## DISCUSSION and CONCLUSION

The *scanMiR* suite provides a fast and flexible way to scan for miRNA binding sites (including potential TDMD and slicing sites), predict transcript repression, and visualize alignments. In addition, we have attempted to make the package user-friendly: *scanMiR* accepts a variety of inputs (e.g. full miRNA sequences, seeds, *KdModels*, etc.) and produces as output standard and well-optimized bioconductor classes, allowing interoperability with other packages. Furthermore, the web interface enables its use by biologists without bioinformatic skills. To our knowledge, this is the first user-friendly tool offering i) transcript-level (rather than gene-level) repression predictions, ii) the prediction of putative TDMD and slicing sites, iii) the ability to scan any custom sequence for high-affinity sites through a web interface, and iv) the possibility to easily extend its functions for any specific species. In addition, *scanMiR* enables the easy and flexible application of McGeary et al. (2019)’s biochemical model of miRNA-mediated repression, significantly outperforming the repression predictions of TargetScan.

An important limitation of the dissociation constant predictions is that they are based on the CNN from McGeary et al. (2019), which was trained on experimental data from only a handful of miRNAs. As a result, predictions for miRNAs whose 12-mer affinity patterns strongly diverge from any of those can be expected to be less accurate. Indeed, whereas the accuracy of scanMiR repression predictions for most CNN-predicted microRNAs is significantly higher than the accuracy obtained with TargetScan score predictions (for some as miR-138 even more than twice as high), for others, it is in fact comparable (miR-217) or even lower (miR-190a) (Fig. 3C). For those miRNAs with a low correlation of predicted and observed repression, limiting the analysis to canonical sites can substantially improve *scanMiR* repression predictions (see miR-190a, Suppl. Fig. 2). This observation is further supported by the fact that imposing very stringent log(K_D_)-cutoffs for these particular miRNAs does not reduce the variance explained, and in one case even improves the correlation (Pearson *r^2^*, Suppl. Fig. 5).

Finally, with time and the accumulation of more experimental data for training, which the *scanMiR* framework is ready to include, we expect those performances to further improve.

Computational predictions of miRNA-mediated repression primarily focusing on the effectiveness of binding sites can only explain actual mRNA changes in cells to a certain degree. In fact, predictions based on the biochemical model can be expected to account for around 50% of the changes in expression attributable to direct miRNA targeting (McGeary et al., 2019). Therefore other critical aspects, such as the accuracy of transcript as well as 3’UTR annotations are likely to improve miRNA target prediction in the future. Studies have suggested that alternative poly-adenylation can have important implications on predicting miRNA-mediated repression in different species and cell types (Nam et al., 2014). Indeed, in the HeLa/HEK cell lines, we observed much lower correlations when using the latest reference human transcriptome instead of the re-assembled line-specific transcriptome (see Methods & Suppl. Fig. 2). Moreover, it has been shown that cis-acting factors, such as RNA-binding proteins (RBPs), can significantly change the binding of miRNAs to certain transcripts (Kedde et al., 2007, 2010; Mubaid et al., 2019). We hence hypothesize that future advances in computational miRNA target prediction may rather be accomplished by the integration of different high-throughput datasets (e.g. RBP-CLIP data or tissue specific smallRNA and mRNA expression data), than by the improvement of affinity predictions of miRNAs to single binding sites.

Finally, an important feature of *scanMiR* is its capability to identify effective non-canonical as well as putative TDMD and slicing sites. To our knowledge, we provide the first systematic screen for potential TDMD sites on linear and circular transcripts highly expressed in neural systems (see also Supplementary Tables 1-2), highlighting several uninvestigated candidates that could play important roles in regulating miRNA abundance. In particular, the scan for enriched circRNA-miRNA pairs in the brain confirmed the exceptional potency of Cdr1as as a highly-expressed miRNA interacting RNA (Guo et al., 2014). In agreement with previous reports of other circRNAs potentially acting as miRNA sponges (Kristensen et al., 2019), a few promising candidates emerged from our analysis which warrant further investigation. We hope that the resources provided here will empower the community to investigate these, and more generally facilitate miRNA research.

## MATERIAL AND METHODS

For detailed information on the individual methods and algorithms, please consult the Bioconductor packages as well as the additional Git repository containing all scripts used in this paper : https://github.com/ETHZ-INS/scanMiR_paper. For a description of the aggregation, external datasets used, their pre-processing and and comparisons, see the Supplementary Methods.

### Encoding of miRNA dissociation constants in *KdModels*

Dissociation constants as obtained from the CNN by McGeary et al. (2019) are stored as a deliberately overfitted linear model, with base K_D_ coefficients for each 1024 partially-matching 8-mers (containing at least four nucleotides (nt) complementary to miRNA positions 1-8, see McGeary et al. (2019)). Subsequently, 8-mer-specific coefficients that are multiplied with a flanking score generated by the flanking di-nucleotides are added to these base 8-mer coefficients. The flanking score is calculated based on the di-nucleotide effects experimentally measured by McGeary et al. (2019). As a result, affinity information for one miRNA can effectively be stored as 2048 integers plus the mature miRNA sequence.

### Sequence scanning

Scanning is performed independently for different miRNAs. When including non-canonical sites, sequence scanning is handled by searching for and extending around four-nucleotide seeds. Coordinates are stored as GRanges (Lawrence et al., 2013), and sequences are extracted using Biostrings. Match type, predicted K_D_, and 3’ local alignment are then estimated for each match. For matches within a given distance from each other, the lowest-affinity match is iteratively excluded until no overlap remains.

Another recent model on Ago-binding affinities fitted on measured miRNA association kinetics of full lengths sequences for miR-21-5p and let-7a-5p suggests (partial) tolerance for G:U wobble binding in the 3’ supplementary region of miRNAs (position 12-17, see also Suppl. Fig. 9). *ScanMiR* therefore includes an option to allow G:U pairing in the 3’ local alignment (Becker et al., 2019). Values for the maximal loop size between the seed and 3’ alignment are by default set to 7 (microRNA) and 9 (target) nucleotides based on recent observations of potential extended effective 3’-supplementary binding of microRNAs (Becker et al., 2019; Sheu-Gruttadauria et al., 2019).

To classify miRNA sites as potentially inducing target-directed miRNA degradation (TDMD) sites, we implemented the following rules adapted from the empirical and theoretical observations of de la Mata et al. (2015), Sheu-Gruttadauria et al. (2019), and Grimson et al. (2007): i) 7mer or 8mer seed-pairing, ii) a 3’-supplementary binding score of at least 6 (to induce the required structural change in the Ago-miRNA complex allowing the ubiquitin ligase Zswim8 to bind (Han et al., 2020; Shi et al., 2020) and releasing the 3’-end of the miRNA (Sheu-Gruttadauria et al., 2019)), iii) a miRNA bulge of at least 2 to avoid central pairing, iv) complementary local alignment of the 3’-end part of the miRNA allowing a maximum of 1 mismatch, v) a maximal target-miRNA bulge difference of 4, and vi) a maximal miRNA bulge of 7.

Efficient miRNA-induced endonucleolytic cleavage (“slicing”) requires central pairing of the miRNA to a target RNA at position 9-11 and is generally thought to be bolstered by complementary pairing of positions 2-15, with single mismatches at specific positions being possibly tolerated (Ameres et al., 2007; Becker et al., 2019). Since a conclusive understanding of miRNA induced target “slicing” remains to be elucidated, we implemented a conservative two-step “slicing” classification that requires: i) complementary binding from positions 2-15 (G:U wobble bindings are allowed outside positions 9-11, though require additional Watson-Crick pairing for each G:U) (denoted “slicing?”), and ii) additional binding from position 16 towards the 3’ end of the miRNA with a maximum of 1 mismatch as well as restricting wobble bindings to miRNA positions beyond the seed (“slicing”).

Beside the scans on the HeLa/HEK transcriptomes, which were performed on the transcriptome reconstructions available in GEO series GSE140217 and GSE140218, the human scans were performed on the GRCh38 ensembl 103 transcriptome (and in addition on the GRCh37 ensembl 75 transcriptome as well as the custom human 3’UTR annotations provided by TargetScan for Suppl. Fig. 2). Unless specified otherwise, the mouse scans were performed on the GRCm38 ensembl 102 transcriptome.

### Web app and indexed random access to large scans

The web interface is a R shiny application which can be deployed through the *scanMiRApp* package. To provide more rapid results, it can optionally use pre-compiled scan and aggregation data. Since the binding site data for all miRNAs and transcripts can amount to a very large size, we implemented the *IndexedFst* S4 class which adds indices to an *fst* file (Klik et al., 2020) to provide very fast out-of-memory access to specific transcripts/miRNAs. The *IndexedFst* class is not limited to storing binding sites and can be used for any data frame or *GenomicRanges* object.

### Software availability

*scanMiR* and its companion packages are available on Bioconductor (see especially https://bioconductor.org/packages/release/bioc/html/scanMiR.html), as well as in the following repositories:

https://github.com/ETHZ-INS/scanMiR
https://github.com/ETHZ-INS/scanMiRData
https://github.com/ETHZ-INS/scanMiRApp

The *scanMIR* web application is available at https://ethz-ins.org/scanMiR/, and can additionally be deployed anywhere using the *scanMiRApp* package.

All processed data and code underlying the figures is available in the paper’s repository, at https://github.com/ETHZ-INS/scanMiR_paper

## SUPPLEMENTARY DATA

Supplementary Methods

Supplementary Figures 1-9

Supplementary Tables 1-4

## ACKNOWLEDGEMENT

We thank Winston Becker for sharing the parameters of their model of Ago2 binding affinity, Kathy Lin for providing insights on how normalizations were calculated for the correlations between observed and predicted repression in McGeary et al. (2019), and Jinyang Zhang for providing the assembled circRNAs from Zhang et al. (2021).

## FUNDING

This work was supported by the Swiss National Science Foundation [SNF_189486 to GS].

## CONFLICT OF INTEREST

The authors declare that they have no conflict of interest.

## Notes

https://github.com/ETHZ-INS/scanMiR

http://ethz-ins.org/scanMiR/

### Competing Interest Statement

The authors have declared no competing interest.

### Summary of Updates

Some of the figures and methods' descriptions were clarified, and TDMD predictions were restricted to miRNAs showing experimental evidence of being TDMD-sensitive.

## REFERENCES

Agarwal, V., Bell, G. W., Nam, J. W., & Bartel, D. P. (2015). Predicting effective microRNA target sites in mammalian mRNAs. eLife, 4, 1–38. https://doi.org/10.7554/eLife.05005

Ameres, S. L., Horwich, M. D., Hung, J.-H., Xu, J., Ghildiyal, M., Weng, Z., & Zamore, P. D. (2010). Target RNA–Directed Trimming and Tailing of Small Silencing RNAs. Science, 328(5985), 1534–1539. https://doi.org/10.1126/science.1187058

Ameres, S. L., Martinez, J., & Schroeder, R. (2007). Molecular Basis for Target RNA Recognition and Cleavage by Human RISC. Cell, 130(1), 101–112. https://doi.org/10.1016/j.cell.2007.04.037

Ameres, S. L., & Zamore, P. D. (2013). Diversifying microRNA sequence and function. Nature Reviews Molecular Cell Biology, 14(8), 475–488. https://doi.org/10.1038/nrm3611

Amin, N. D., Bai, G., Klug, J. R., Bonanomi, D., Pankratz, M. T., Gifford, W. D., Hinckley, C. A., Sternfeld, M. J., Driscoll, S. P., Dominguez, B., Lee, K. F., Jin, X., & Pfaff, S. L. (2015). Loss of motoneuron-specific microRNA-218 causes systemic neuromuscular failure. Science, 350(6267), 1525–1529. https://doi.org/10.1126/science.aad2509

Bartel, D. P. (2004). MicroRNAs: Genomics, Biogenesis, Mechanism, and Function. Cell, 116(2), 281–297. https://doi.org/10.1016/S0092-8674(04)00045-5

Bartel, D. P. (2009). MicroRNAs: Target Recognition and Regulatory Functions. Cell, 136(2), 215–233. https://doi.org/10.1016/j.cell.2009.01.002

Bartel, D. P. (2018). Metazoan MicroRNAs. Cell, 173(1), 20–51. https://doi.org/10.1016/j.cell.2018.03.006

Becker, W. R., Ober-Reynolds, B., Jouravleva, K., Jolly, S. M., Zamore, P. D., & Greenleaf, W. J. (2019). High-Throughput Analysis Reveals Rules for Target RNA Binding and Cleavage by AGO2. Molecular Cell, 1–15. https://doi.org/10.1016/j.molcel.2019.06.012

Bitetti, A., Mallory, A. C., Golini, E., Carrieri, C., Carreño Gutiérrez, H., Perlas, E., Pérez-Rico, Y. A., Tocchini-Valentini, G. P., Enright, A. J., Norton, W. H. J., Mandillo, S., O’Carroll, D., & Shkumatava, A. (2018). MicroRNA degradation by a conserved target RNA regulates animal behavior. Nature Structural & Molecular Biology, 25(3), 244–251. https://doi.org/10.1038/s41594-018-0032-x

Blankenberg, D., Taylor, J., Nekrutenko, A., & The Galaxy Team. (2011). Making whole genome multiple alignments usable for biologists. Bioinformatics, 27(17), 2426–2428. https://doi.org/10.1093/bioinformatics/btr398

Carthew, R. W., & Sontheimer, E. J. (2009). Origins and Mechanisms of miRNAs and siRNAs. Cell, 136(4), 642–655. https://doi.org/10.1016/j.cell.2009.01.035

Chi, S. W., Hannon, G. J., & Darnell, R. B. (2012). An alternative mode of microRNA target recognition. Nature Structural and Molecular Biology, 19(3), 321–327. https://doi.org/10.1038/nsmb.2230

Chiang, H. R., Schoenfeld, L. W., Ruby, J. G., Auyeung, V. C., Spies, N., Baek, D., Johnston, W. K., Russ, C., Luo, S., Babiarz, J. E., Blelloch, R., Schroth, G. P., Nusbaum, C., & Bartel, D. P. (2010). Mammalian microRNAs: Experimental evaluation of novel and previously annotated genes. Genes & Development, 24(10), 992–1009. https://doi.org/10.1101/gad.1884710

de la Mata, M., Gaidatzis, D., Vitanescu, M., Stadler, M. B., Wentzel, C., Scheiffele, P., Filipowicz, W., & Grosshans, H. (2015). Potent degradation of neuronal miRNAs induced by highly complementary targets. EMBO reports, 16(4), 500–511. https://doi.org/10.15252/embr.201540078

Denzler, R., McGeary, S. E., Title, A. C., Agarwal, V., Bartel, D. P., & Stoffel, M. (2016). Impact of MicroRNA Levels, Target-Site Complementarity, and Cooperativity on Competing Endogenous RNA-Regulated Gene Expression. Molecular Cell, 64(3), 565–579. https://doi.org/10.1016/j.molcel.2016.09.027

Eichhorn, S. W., Guo, H., McGeary, S. E., Rodriguez-Mias, R. A., Shin, C., Baek, D., Hsu, S., Ghoshal, K., Villén, J., & Bartel, D. P. (2014). MRNA Destabilization Is the Dominant Effect of Mammalian MicroRNAs by the Time Substantial Repression Ensues. Molecular Cell, 56(1), 104–115. https://doi.org/10.1016/j.molcel.2014.08.028

Fuchs Wightman, F., Giono, L. E., Fededa, J. P., & de la Mata, M. (2018). Target RNAs Strike Back on MicroRNAs. Frontiers in Genetics, 9. https://www.frontiersin.org/article/10.3389/fgene.2018.00435

Garcia, D. M., Baek, D., Shin, C., Bell, G. W., Grimson, A., & Bartel, D. P. (2011). Weak seed-pairing stability and high target-site abundance decrease the proficiency of lsy-6 and other microRNAs. Nature Structural & Molecular Biology, 18(10), 1139–1146. https://doi.org/10.1038/nsmb.2115

Griffiths-Jones, S., Grocock, R. J., van Dongen, S., Bateman, A., & Enright, A. J. (2006). miRBase: MicroRNA sequences, targets and gene nomenclature. Nucleic Acids Research, 34(suppl_1), D140–D144. https://doi.org/10.1093/nar/gkj112

Grimson, A., Farh, K. K. H., Johnston, W. K., Garrett-Engele, P., Lim, L. P., & Bartel, D. P. (2007). MicroRNA Targeting Specificity in Mammals: Determinants beyond Seed Pairing. Molecular Cell, 27(1), 91–105. https://doi.org/10.1016/j.molcel.2007.06.017

Gumienny, R., & Zavolan, M. (2015). Accurate transcriptome-wide prediction of microRNA targets and small interfering RNA off-targets with MIRZA-G. Nucleic Acids Research, 43(3), 1380–1391. https://doi.org/10.1093/nar/gkv050

Guo, J. U., Agarwal, V., Guo, H., & Bartel, D. P. (2014). Expanded identification and characterization of mammalian circular RNAs. Genome Biology, 15(7), 409. https://doi.org/10.1186/s13059-014-0409-z

Han, J., LaVigne, C. A., Jones, B. T., Zhang, H., Gillett, F., & Mendell, J. T. (2020). A ubiquitin ligase mediates target-directed microRNA decay independently of tailing and trimming. Science, 370(6523). https://doi.org/10.1126/science.abc9546

Hansen, T. B., Wiklund, E. D., Bramsen, J. B., Villadsen, S. B., Statham, A. L., Clark, S. J., & Kjems, J. (2011). MiRNA-dependent gene silencing involving Ago2-mediated cleavage of a circular antisense RNA. The EMBO Journal, 30(21), 4414–4422. https://doi.org/10.1038/emboj.2011.359

Hausser, J., Landthaler, M., Jaskiewicz, L., Gaidatzis, D., & Zavolan, M. (2009). Relative contribution of sequence and structure features to the mRNA binding of Argonaute/EIF2C–miRNA complexes and the degradation of miRNA targets. Genome Research, 19(11), 2009–2020. https://doi.org/10.1101/gr.091181.109

Jonas, S., & Izaurralde, E. (2015). Towards a molecular understanding of microRNA-mediated gene silencing. Nature Reviews Genetics, 16(7), 421–433. https://doi.org/10.1038/nrg3965

Kedde, M., Strasser, M. J., Boldajipour, B., Vrielink, J. A. F. O., Slanchev, K., le Sage, C., Nagel, R., Voorhoeve, P. M., van Duijse, J., Ørom, U. A., Lund, A. H. H., Perrakis, A., Raz, E., & Agami, R. (2007). RNA-Binding Protein Dnd1 Inhibits MicroRNA Access to Target mRNA. Cell, 131(7), 1273–1286. https://doi.org/10.1016/j.cell.2007.11.034

Kedde, M., van Kouwenhove, M., Zwart, W., Oude Vrielink, J. A. F., Elkon, R., & Agami, R. (2010). A Pumilio-induced RNA structure switch in p27-3’UTR controls miR-221 and miR-222 accessibility. Nature Cell Biology, 12(10), 1014–1020. https://doi.org/10.1038/ncb2105

Kern, F., Backes, C., Hirsch, P., Fehlmann, T., Hart, M., Meese, E., & Keller, A. (2020). What’s the target: Understanding two decades of in silico microRNA-target prediction. Briefings in Bioinformatics, 21(6), 1999–2010. https://doi.org/10.1093/bib/bbz111

Kleaveland, B., CY, S., Stefano, J., & DP, B. (2018). A Network of Noncoding Regulatory RNAs Acts in the Mammalian Brain. LID - S0092-8674(18)30634-2 [pii] LID - 10.1016/j.cell.2018.05.022 [doi]. 1097–4172 (Electronic). https://doi.org/10.1101/279687

Klik, M., LZ4), Y C. (Yann C. is author of the bundled L. and Z. code and copyright holder of, Facebook, & code), I. (Bundled Z. (2020). fst: Lightning Fast Serialization of Data Frames (Version 0.9.4) [Computer software]. https://CRAN.R-project.org/package=fst

Krek, A., Grün, D., Poy, M. N., Wolf, R., Rosenberg, L., Epstein, E. J., MacMenamin, P., da Piedade, I., Gunsalus, K. C., Stoffel, M., & Rajewsky, N. (2005). Combinatorial microRNA target predictions. Nature Genetics, 37(5), 495–500. https://doi.org/10.1038/ng1536

Kristensen, L. S., Andersen, M. S., Stagsted, L. V. W., Ebbesen, K. K., Hansen, T. B., & Kjems, J. (2019). The biogenesis, biology and characterization of circular RNAs. Nature Reviews Genetics. https://doi.org/10.1038/s41576-019-0158-7

Lawrence, M., Huber, W., Pagès, H., Aboyoun, P., Carlson, M., Gentleman, R., Morgan, M. T., & Carey, V. J. (2013). Software for Computing and Annotating Genomic Ranges. PLOS Computational Biology, 9(8), e1003118. https://doi.org/10.1371/journal.pcbi.1003118

Liu, W., & Wang, X. (2019). Prediction of functional microRNA targets by integrative modeling of microRNA binding and target expression data. Genome Biology, 20(1), 18. https://doi.org/10.1186/s13059-019-1629-z

McGeary, S. E., Bisaria, N., & Bartel, D. P. (2021). Pairing to the microRNA 3’region occurs through two alternative binding modes, with affinity shaped by nucleotide identity as well as pairing position. BioRxiv, 2021.04.13.439700. https://doi.org/10.1101/2021.04.13.439700

McGeary, S. E., Lin, K. S., Shi, C. Y., Pham, T. M., Bisaria, N., Kelley, G. M., & Bartel, D. P. (2019). The biochemical basis of microRNA targeting efficacy. Science, 366(6472). https://doi.org/10.1126/science.aav1741

Mubaid, S., Ma, J. F., Omer, A., Ashour, K., Lian, X. J., Sanchez, B. J., Robinson, S., Cammas, A., Dormoy-Raclet, V., Di Marco, S., Chittur, S. V., Tenenbaum, S. A., & Gallouzi, I.-E. (2019). HuR counteracts miR-330 to promote STAT3 translation during inflammation-induced muscle wasting. Proceedings of the National Academy of Sciences, 201905172–201905172. https://doi.org/10.1073/pnas.1905172116

Nam, J.-W., Rissland, O. S., Koppstein, D., Abreu-Goodger, C., Jan, C. H., Agarwal, V., Yildirim, M. A., Rodriguez, A., & Bartel, D. P. (2014). Global Analyses of the Effect of Different Cellular Contexts on MicroRNA Targeting. Molecular Cell, 53(6), 1031–1043. https://doi.org/10.1016/j.molcel.2014.02.013

Nielsen, C. B., Shomron, N., Sandberg, R., Hornstein, E., Kitzman, J., & Burge, C. B. (2007). Determinants of targeting by endogenous and exogenous microRNAs and siRNAs. RNA, 13(11), 1894–1910. https://doi.org/10.1261/rna.768207

Patro, R., Duggal, G., Love, M. I., Irizarry, R. A., & Kingsford, C. (2017). Salmon provides fast and bias-aware quantification of transcript expression. Nature Methods, 14(4), 417–419. https://doi.org/10.1038/nmeth.4197

Reczko, M., Maragkakis, M., Alexiou, P., Grosse, I., & Hatzigeorgiou, A. G. (2012). Functional microRNA targets in protein coding sequences. Bioinformatics, 28(6), 771–776. https://doi.org/10.1093/bioinformatics/bts043

Robinson, M. D., McCarthy, D. J., & Smyth, G. K. (2010). edgeR: A Bioconductor package for differential expression analysis of digital gene expression data. Bioinformatics, 26(1), 139–140. https://doi.org/10.1093/bioinformatics/btp616

Sætrom, P., Heale, B. S. E., Snøve, O., Aagaard, L., Alluin, J., & Rossi, J. J. (2007). Distance constraints between microRNA target sites dictate efficacy and cooperativity. Nucleic Acids Research, 35(7), 2333–2342. https://doi.org/10.1093/nar/gkm133

Salomon, W. E., Jolly, S. M., Moore, M. J., Zamore, P. D., & Serebrov, V. (2015). Single-Molecule Imaging Reveals that Argonaute Reshapes the Binding Properties of Its Nucleic Acid Guides. Cell, 162(1), 84–95. https://doi.org/10.1016/j.cell.2015.06.029

Schirle, N. T., Sheu-Gruttadauria, J., & MacRae, I. J. (2014). Structural basis for microRNA targeting. Science, 346(6209), 608–613. https://doi.org/10.1126/science.1258040

Sheu-Gruttadauria, J., Pawlica, P., Klum, S. M., Wang, S., Yario, T. A., Schirle Oakdale, N. T., Steitz, J. A., & MacRae, I. J. (2019). Structural Basis for Target-Directed MicroRNA Degradation. Molecular Cell, 75(6), 1243–1255.e7. https://doi.org/10.1016/j.molcel.2019.06.019

Sheu-Gruttadauria, J., Xiao, Y., Gebert, L. F., & MacRae, I. J. (2019). Beyond the seed: Structural basis for supplementary micro RNA targeting by human Argonaute2. The EMBO Journal, 38(13), 1–14. https://doi.org/10.15252/embj.2018101153

Shi, C. Y., Kingston, E., Kleaveland, B., Lin, D. H., Stubna, M. W., & Bartel, D. P. (2020). The ZSWIM8 ubiquitin ligase mediates target-directed microRNA degradation. Science, 21(1), 1–9. https://doi.org/10.1126/science.abc9359

Wang, Y., Soneson, C., Malinowska, A. L., Laski, A., Ghosh, S., Kanitz, A., Gebert, L. F. R., Robinson, M. D., & Hall, J. (2021). MiR-CLIP reveals iso-miR selective regulation in the miR-124 targetome. Nucleic Acids Research, 49(1), 25–37. https://doi.org/10.1093/nar/gkaa1117

Whipple, A. J., Breton-provencher, V., Jacobs, H. N., Chitta, U. K., Sur, M., Sharp, P. A., Whipple, A. J., Breton-provencher, V., Jacobs, H. N., Chitta, U. K., & Sur, M. (2020). Imprinted Maternally Expressed microRNAs Antagonize Paternally Driven Gene Programs in Article Imprinted Maternally Expressed microRNAs Antagonize Paternally Driven Gene Programs in Neurons. Molecular Cell, 1–11. https://doi.org/10.1016/j.molcel.2020.01.020

Yekta, S., Shih, I. -hung, & Bartel, D. P. (2004). MicroRNA-Directed Cleavage of HOXB8 mRNA. Science. https://doi.org/10.1126/science.1097434

Zhang, J., Hou, L., Zuo, Z., Ji, P., Zhang, X., Xue, Y., & Zhao, F. (2021). Comprehensive profiling of circular RNAs with nanopore sequencing and CIRI-long. Nature Biotechnology, 39(7), 836–845. https://doi.org/10.1038/s41587-021-00842-6

