## Supplementary material for "scanMiR: a biochemically-based toolkit for versatile and efficient microRNA target prediction": Suppl. Fig.

Supplementary Figures

Michael Soutschek Fridolin Gross Gerhard Schratt Pierre-Luc Germain

22 January, 2022

### Supplementary Figure S1

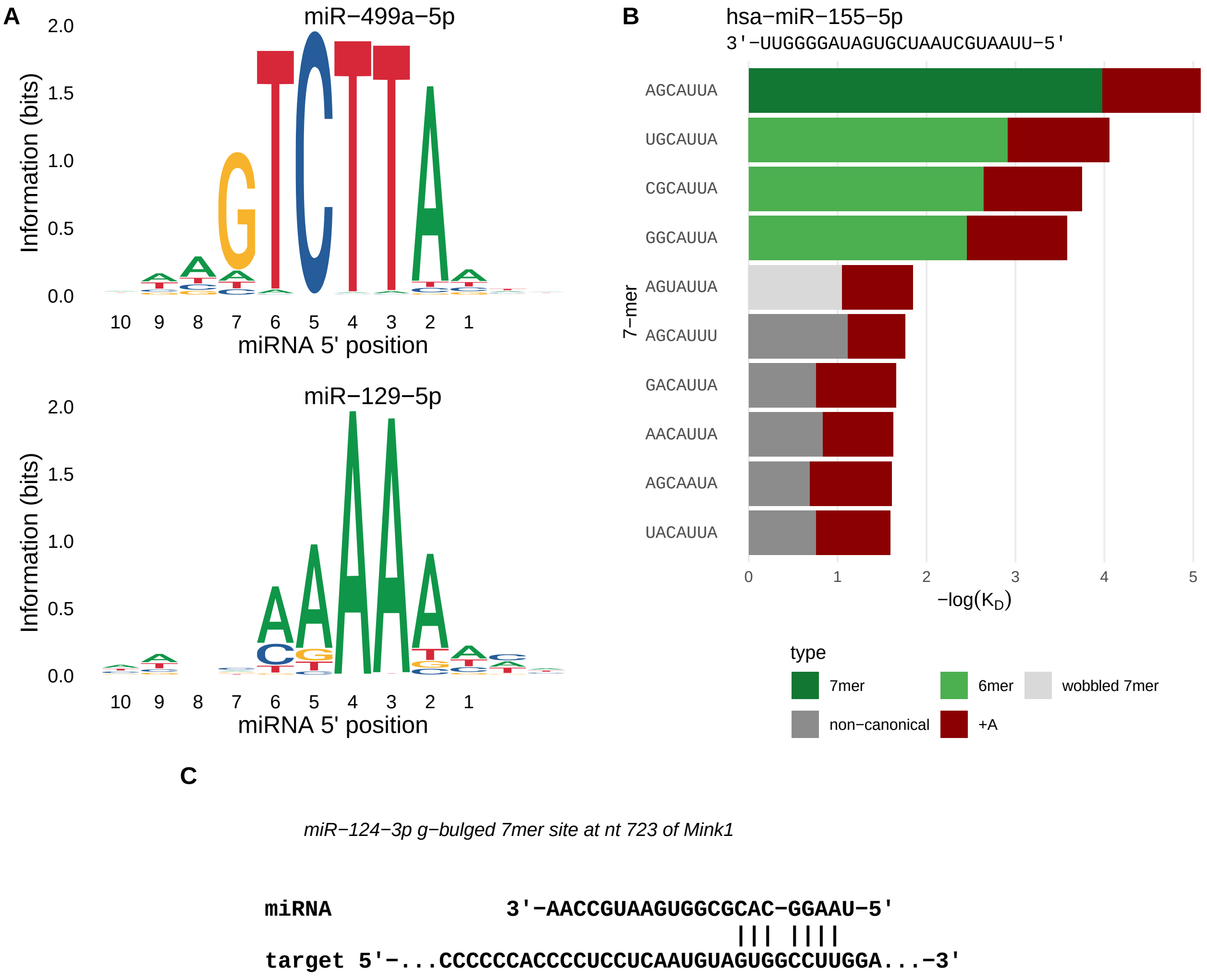


**Supplementary Figure S1**

**ScanMiR plotting functions provide insights into miRNA biology. A:** Nucleotide information content plots of miR-499a-5p and miR-129-5p. The information content of each nucleotide is calculated by its occurrence in the top ten seed complements with the lowest dissociation constants. **B:** Affinity plot of miR-155-5p showing a ‘wobbled 7mer’ as highest ranking non-canonical seed complement. **C:** scanMiR flags g-bulged miRNA binding sites in scan results and correctly represents target alignments of those sites (here shown for the experimentally validated miR-124-3p binding site on Mink1).

### Supplementary Figure S2

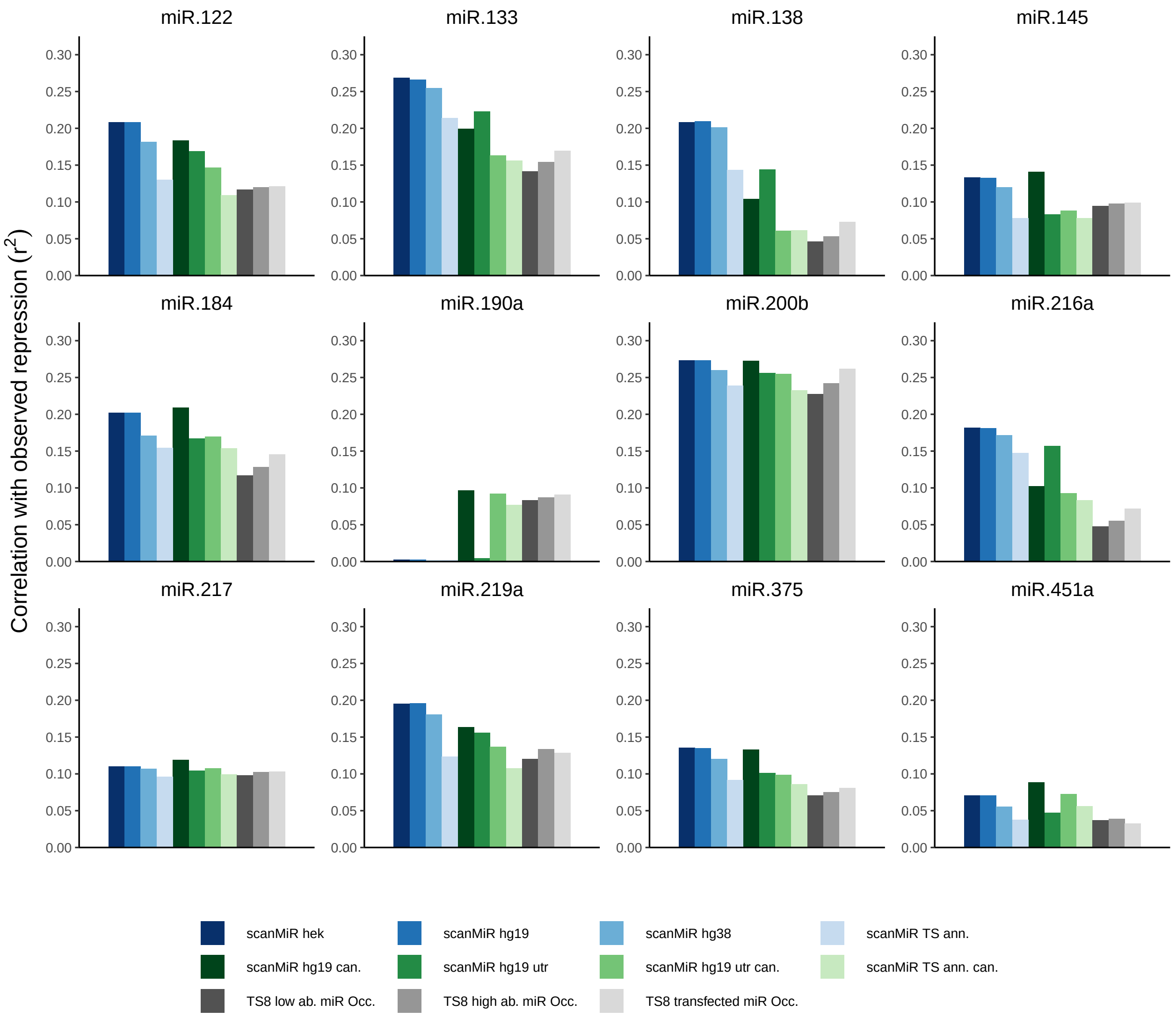


**Supplementary Figure S2**

**Comparison of scanMiR repression predictions versus TargetScan8 occupancy scores.** Pearson correlations of scanMiR repression predictions and TargetScan8 (TS8) occupancy scores with measured mRNA changes following miRNA mimic transfections in HEK cells (McGeary et al., 2019). The first four bars for each miRNA indicate the correlations of default scanMiR repression predictions obtained with different genome annotations (hek = Custom HEK cell annotation from McGeary et al. (2019), hg19 = GRCh37, hg38 = GRCh38 & TS ann. = Custom human 3'UTR annotation obtained from TargetScan8). To comprehensively compare scanMiR repression scores to TargetScan8 occupancy scores, we further included analyses considering either exclusively canonical sites ("can."), only sites in the 3'UTR ("utr") or a combination of both. The last three bars show the Pearson correlations of occupancy scores (for low and high abundant miRNAs as well as transfected miRNAs) provided in the latest TargetScan update (version 8) with the same measured mRNA logFC. All scanMiR repression predictions are calculated with the globally optimized a-value (see Fig. 3C).

### Supplementary Figure S3

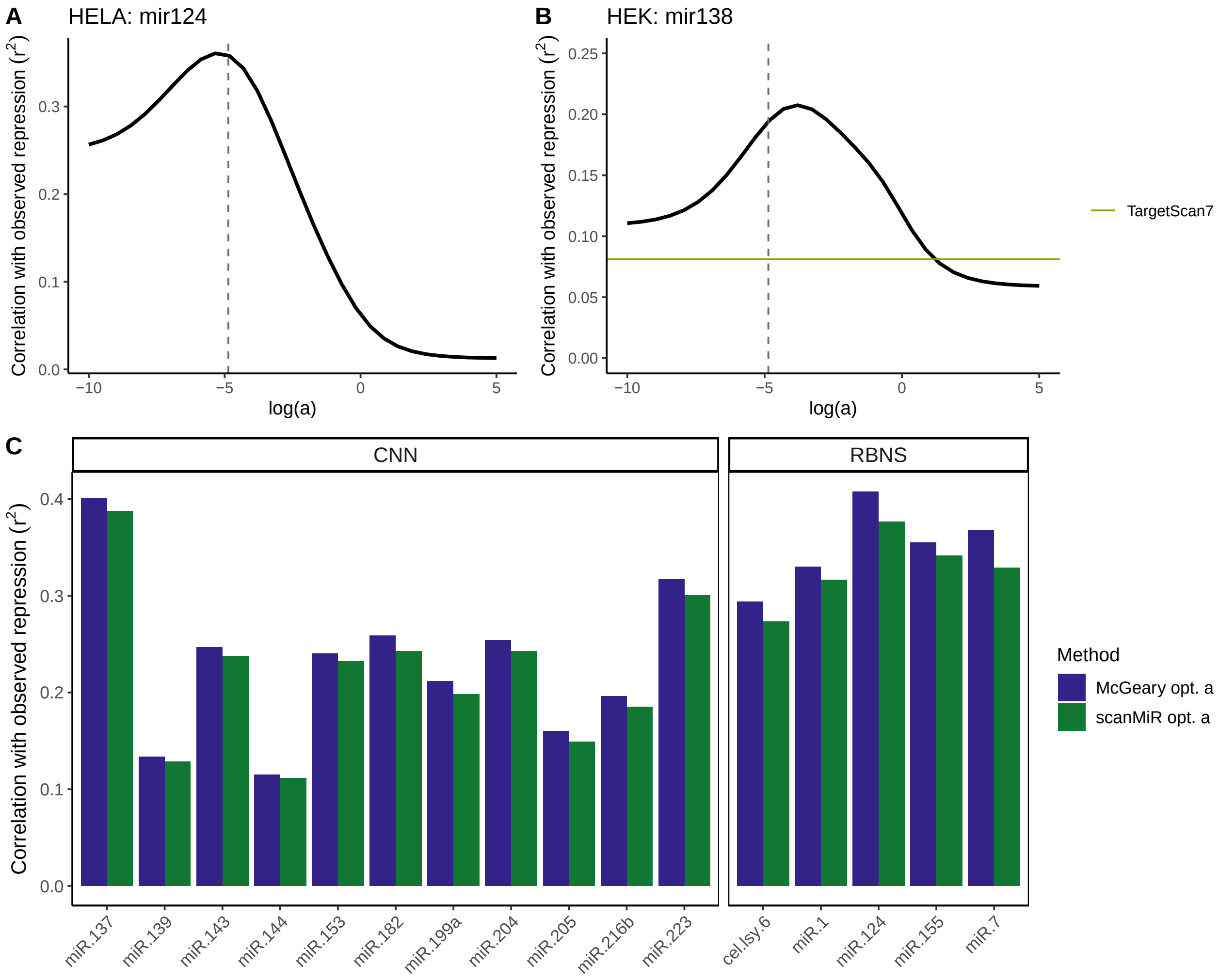


**Supplementary Figure S3**

**Pearson correlations of scanMiR repression predictions to measured changes in HeLa cells. A-B:** Deviations of *a* from the optimal value still lead to high correlations of scanMiR repression predictions with measured mRNA logFC in cells upon miRNA mimic transfections, outperforming TargetScan7 predictions over several orders of magnitudes (**B**). **C:** The correlations of predicted repression to observed foldchange upon miRNA-mimic transfection obtained with scanMiR is comparable to those obtained from the original CNN-values from McGeary et al. (2019). miRNAs used in RBNS experiments are displayed on the right side.

### Supplementary Figure S4

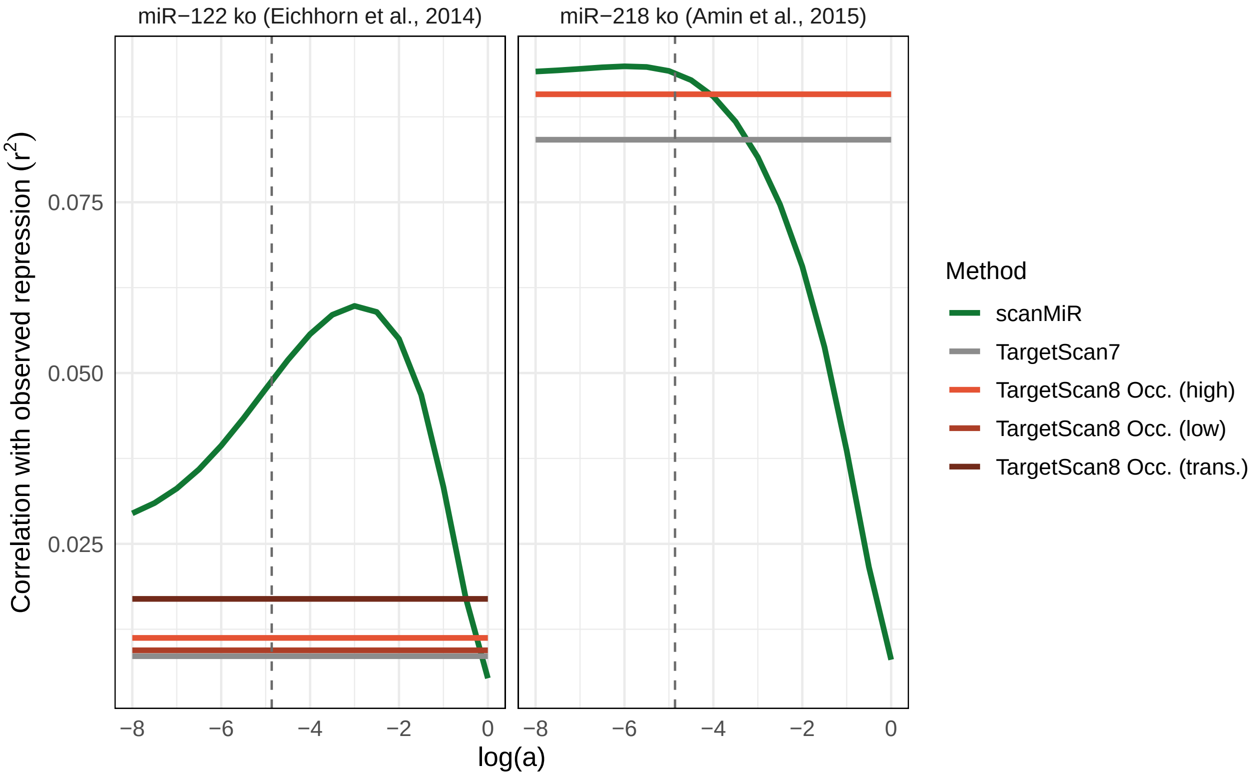


**Supplementary Figure S4**

**ScanMiR repression predictions outperform TargetScan predictions in different mouse miRNA knockout datasets.** Plots displaying the variance explained ($r^2$) of scanMiR and TargetScan repression predictions with mRNA logFC obtained by RNA-sequencing upon knockout of miR-122 (Eichhorn et al., 2014) or miR-218 (Amin et al., 2015). TargetScan8 provides predicted miRNA occupancy scores for highly & lowly expressed miRNAs as well as for transfected (=trans.) miRNAs, whereas TargetScan7 results are based on the context++ score. In the miR-218 panel, only one occupancy score is depicted since the other two are at the same level. The globally optimized log(a)-value used by default in scanMiR is indicated with a dashed grey line.

### Supplementary Figure S5

A

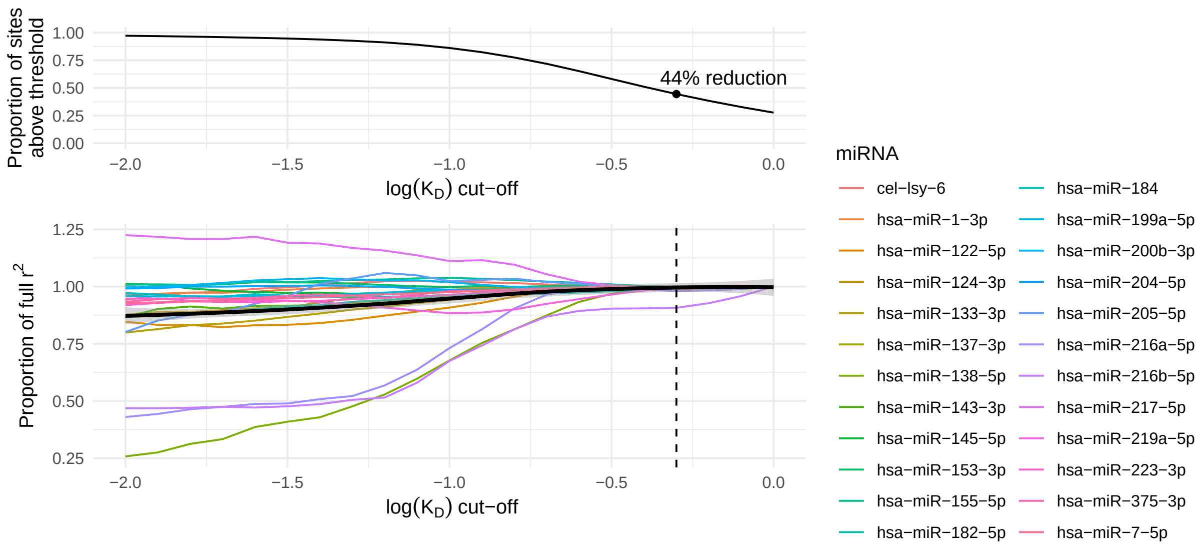


B

| miRNA | Cut−off 97.5% | Cut−off 99% |
|---|---|---|
| hsa−miR−122−5p | −0.7 | −0.6 |
| hsa−miR−133−3p | −0.7 | −0.4 |
| hsa−miR−138−5p | −0.4 | −0.3 |
| hsa−miR−145−5p | −2.0 | −2.0 |
| hsa−miR−184 | −0.5 | −0.3 |
| hsa−miR−190a−5p | −2.0 | −2.0 |
| hsa−miR−200b−3p | −1.7 | −1.6 |
| hsa−miR−216a−5p | −0.6 | −0.5 |
| hsa−miR−217−5p | −2.0 | −2.0 |
| hsa−miR−219a−5p | −0.4 | −0.3 |
| hsa−miR−375−3p | −1.0 | −0.6 |
| hsa−miR−451a | −2.0 | −2.0 |

C

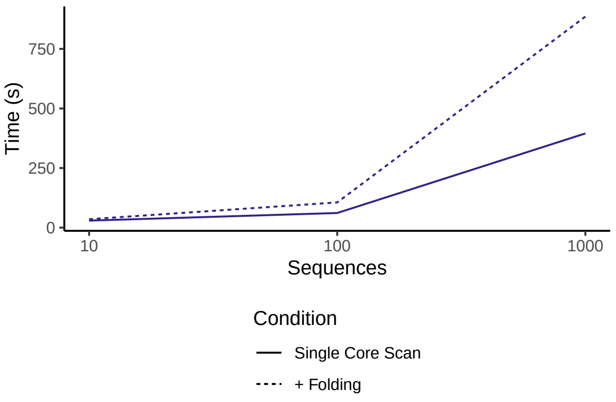


#### Supplementary Figure S5

**Computational optimizations in scanMiR - Preliminary. A:** Proportion of maximal Pearson correlation values of predicted and measured miRNA repression depending on different maximal $K_D$-values (“cutoff”) for all miRNAs transfected in HEK & HeLa with an $r^2$-value of at least 10% from the study of McGeary and colleagues (2019). MiR-216b-5p is the only miRNA displayed with a reduction of more than 2.5% of its maximal $r^2$-value with a cutoff of -0.3. The black line indicates the smoothed mean. Potential reductions in site numbers across the spectrum of indicated cutoffs are shown in the upper panel. **B:** The maximum $log(K_D)$ cutoff-values for the 12 miRNAs used in HEK-transfection experiments (McGeary et al., 2019) which cause a diminution of the correlation by only 2.5% (= Cut-off 97.5%) or 1% (= Cut-off 99%), respectively. For individual miRNAs, there were pronounced variations in the cutoff beyond which the addition of binding sites with a lower affinity did not substantially improve the correlation between predicted and observed repression. **C:** Run times of the scanning algorithm of McGeary et al. (2019) with and without folding of the transcripts.

#### Supplementary Figure S6

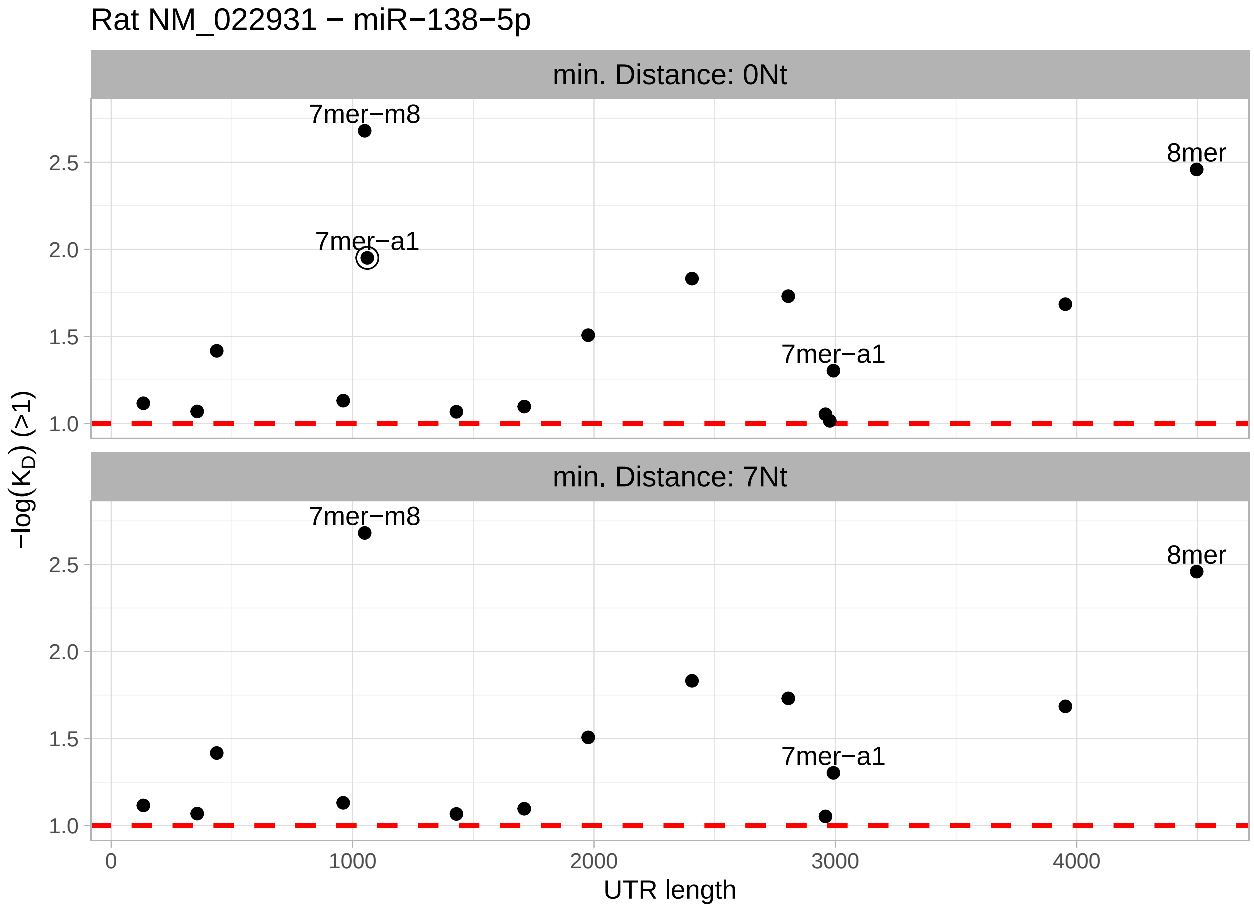


**Supplementary Figure S6**

**Removal of overlapping sites in scanMiR.** Users can specify a minimum allowed distance between binding sites of the same miRNA. Within that distance, scanMiR automatically keeps only the site with the highest affinity, as shown here for miR-138-5p binding sites on a rat Rims3 transcript. An example site removed due to a nearby higher-affinity site is highlighted.

### Supplementary Figure S7

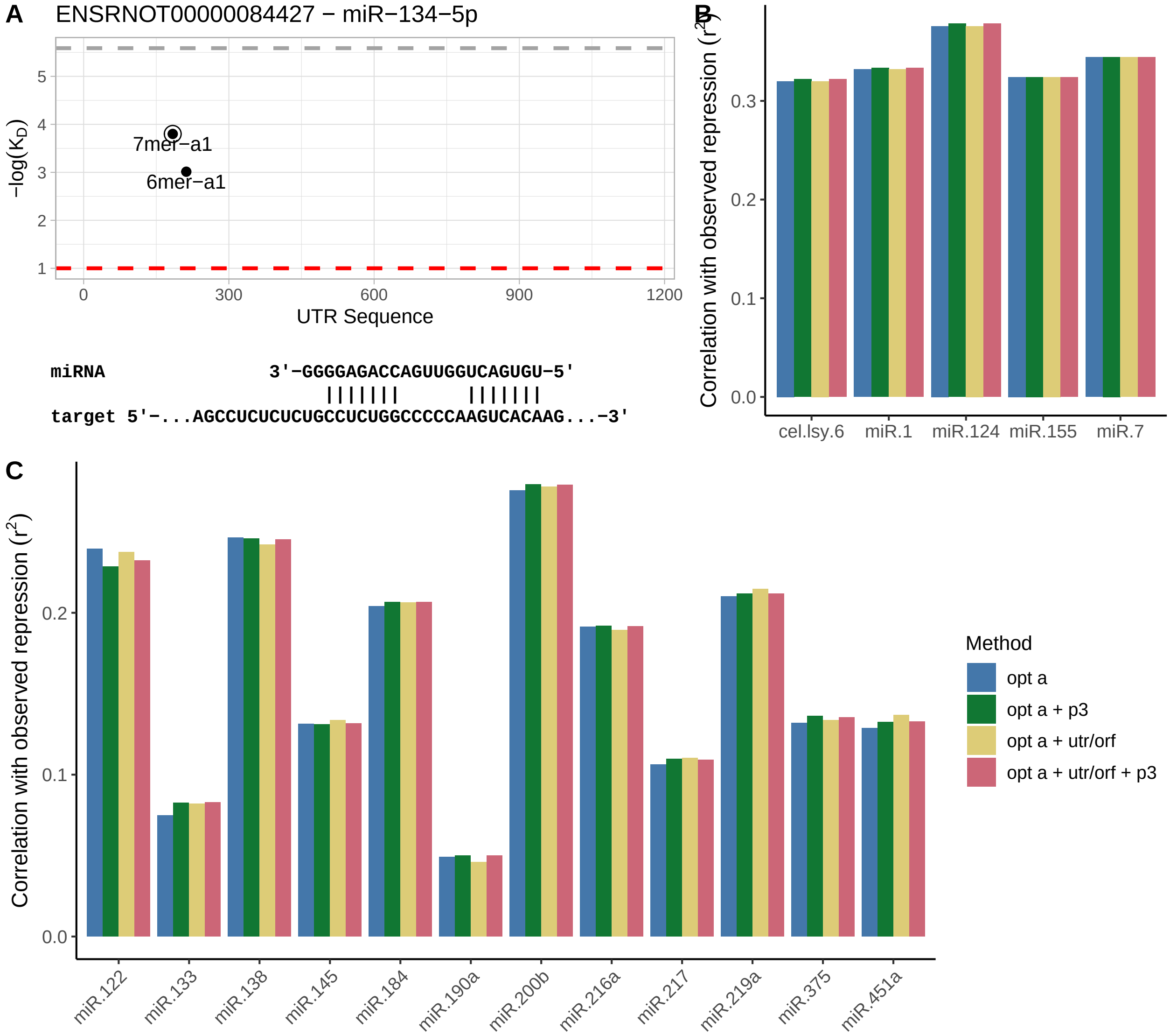


**Supplementary Figure S7**

**The 3'-supplementary score as well as length coefficients implemented in scanMiR slightly improve repression prediction. A:** Example miRNA binding site including extensive 3'-supplementary pairing. **B-C:** Pearson correlation values of observed and predicted repression in HeLA- (**B**) and HEK- (**C**) cells, including 3'-supplementary score as well as coefficients taking into account the open reading frame (ORF) and 3'UTR lengths of transcripts.

### Supplementary Figure S8

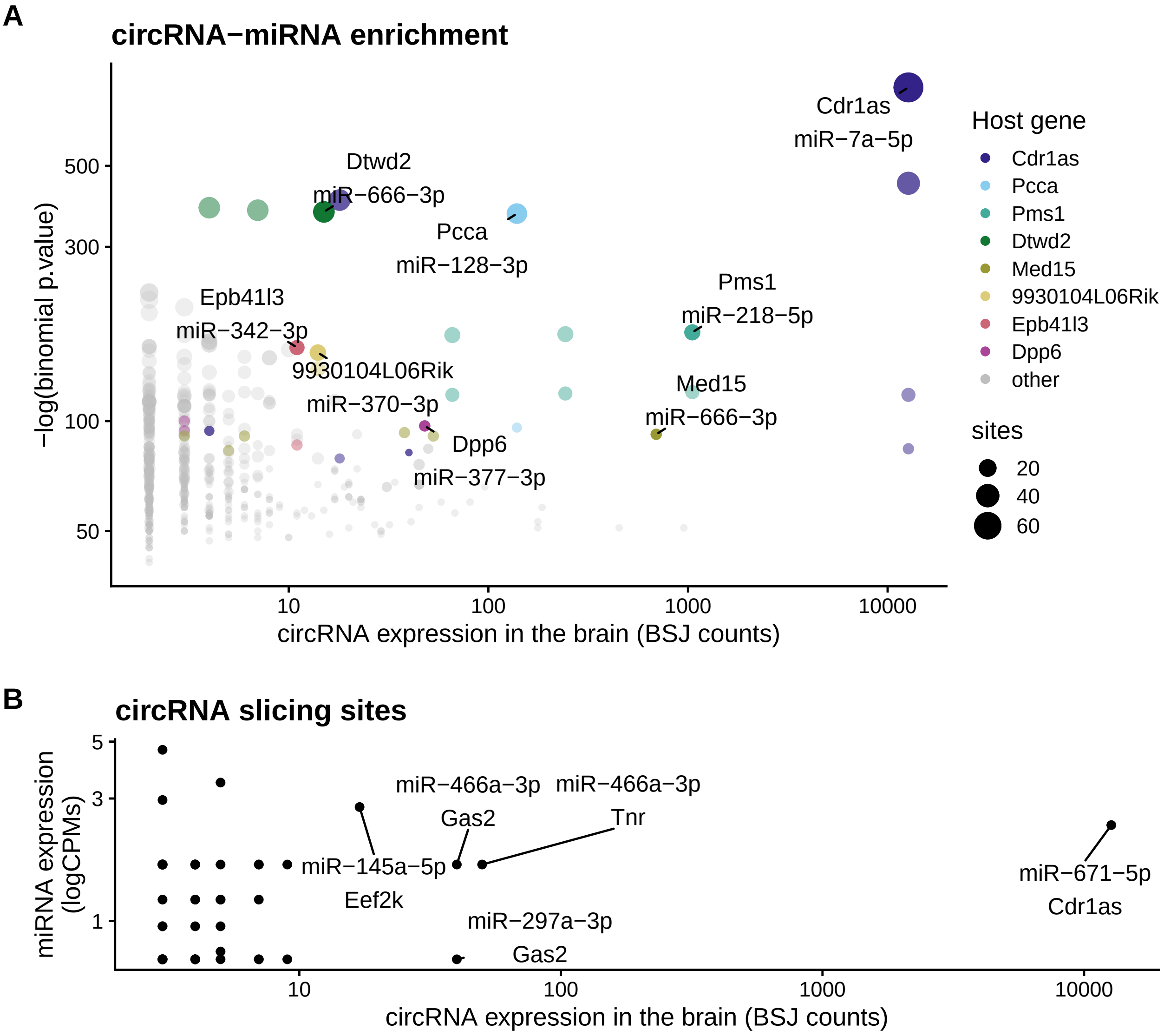


**Supplementary Figure S8**

**Prediction of key miRNA binding sites on circRNAs. A:** Abundant circRNAs with enrichment for specific miRNA binding sites. Each dot represents a miRNA and circular RNA isoform pair, with only the most significant pair for interesting candidate genes being labeled. The y-axis indicates the significance of the circRNA-miRNA enrichment. Only miRNAs with at least 100 reads and circRNAs overlapping genes were considered. **B:** Prediction of circRNA slicing sites. The by far strongest signal comes from the Cdr1as circRNA.

### Supplementary Figure S9

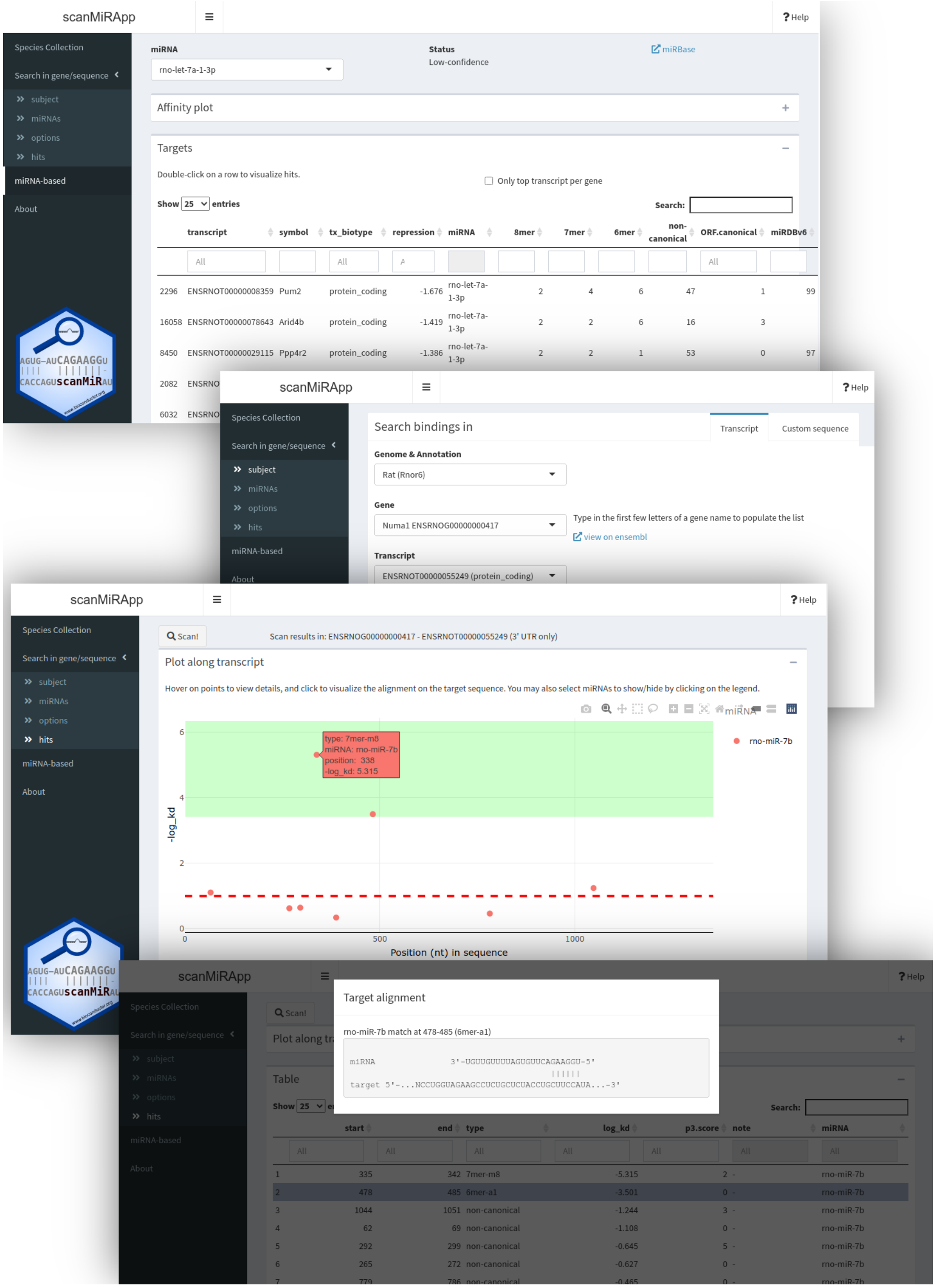


#### Supplementary Figure S9

**Screenshots showing some of the functionalities of the scanMiR web application.** The live app can be visited at https://ethz-ins.org/scanMiR/

#### Supplementary Figure S10

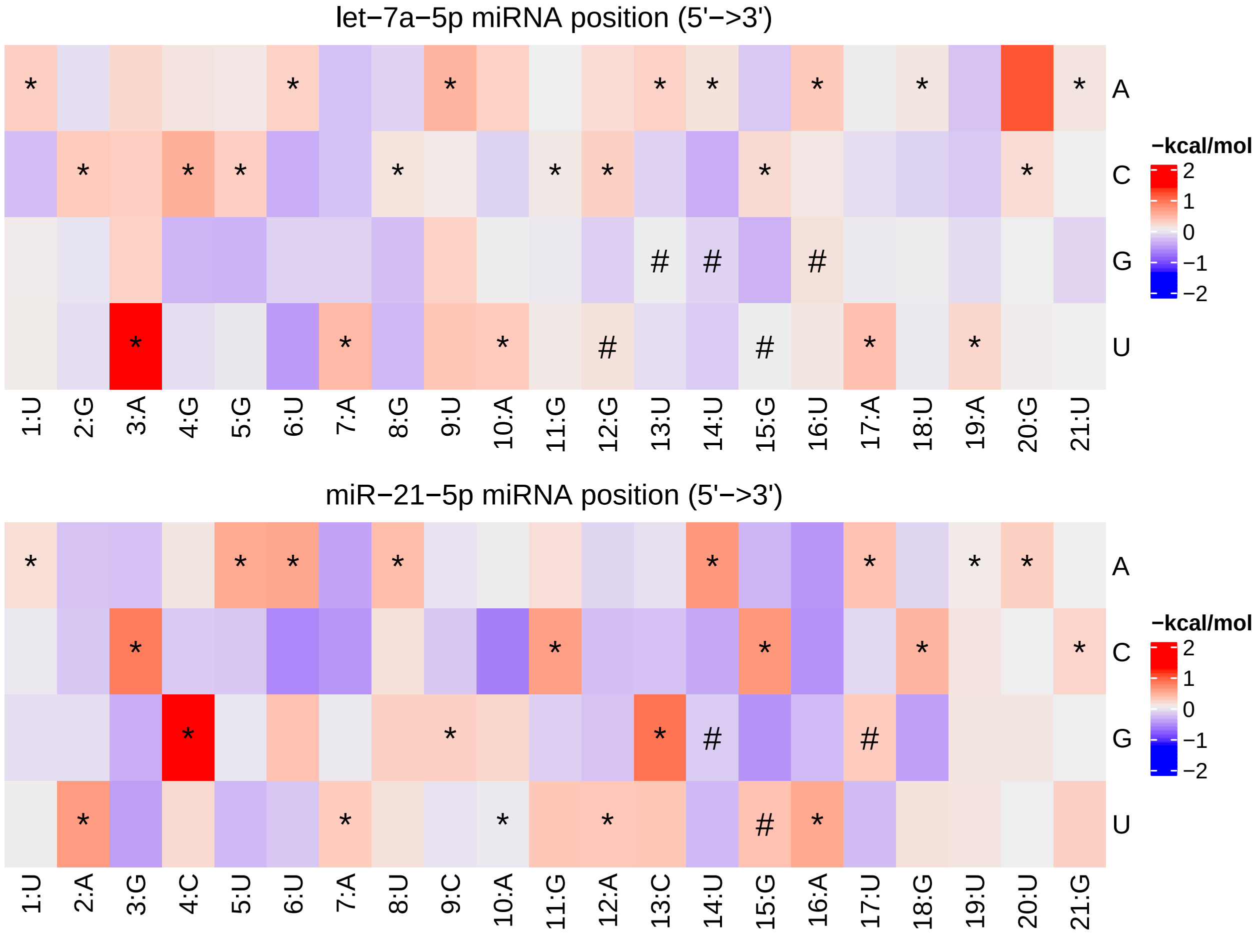


**Supplementary Figure S10**

**Modelled Ago-binding affinities for two full length miRNA sequences.** Becker et al. (2019) recently measured association kinetics of Ago2 loaded with miRNA let-7a-5p and miR-21-5p to ca. 20'000 RNA targets each. Based on this data, they subsequently modelled positional Ago-Binding affinities for both of the miRNA-sequences. Plotted are these predicted affinity values towards each nucleotide of the miRNA-sequences (depicted on the x-axis). Asterics indicate the Watson-Crick pairing at each position and hashes potential G:U wobble bindings at positions 12-17.
