## Supplementary Methods for "scanMiR: a biochemically-based toolkit for versatile and efficient microRNA target prediction"

Michael Soutschek Fridolin Gross Gerhard Schratt Pierre-Luc Germain

22 Januar, 2022

#### Aggregation into predicted transcript repression

The aggregation of multiple binding sites into predicted transcript repression is done mostly according to the model by McGeary, Lin et al. (2019). Briefly, multiple miRNA binding sites are assumed to have an additive effect on the transcript's decay rate that is proportional to their occupancy. The occupancy of AGO-bound miRNA $g$ on a mRNA $m$ with $p$ binding sites in the open reading frame (ORF) and $q$ in the 3' untranslated region (UTR) is given by the following equation:

$$N_{m,g} = \sum_{i=1}^{p} \left( \frac{a_g}{a_g + c_{\mathrm{ORF}} K_{d,i}^{\mathrm{ORF}}} \right) + \sum_{j=1}^{q} \left( \frac{a_g}{a_g + K_{d,j}^{\mathrm{3'UTR}}} \right)$$

where $a$ is the relative concentration of unbound AGO-miRNA complexes, and $c$ is the penalty factor for sites that are found within the ORF. scanMiR also includes a coefficient $e$ accounting for the effect of the 3' alignment:

$$N_{m,g} = \sum_{i=1}^{p} \left( \frac{a_g}{a_g + e_i c_{\mathrm{ORF}} K_{d,i}^{\mathrm{ORF}}} \right) + \sum_{j=1}^{q} \left( \frac{a_g}{a_g + e_j K_{d,j}^{\mathrm{3'UTR}}} \right)$$

where $e$ is the exponential of the product of the 3' alignment score and a global parameter ($p3$). The 3' alignment score roughly corresponds to the number of matched nucleotides (additionally penalizing gaps), and it is by default set to 0 if the number of matches is below 3 (adapted from the observations in Grimson et al., 2007) and capped to a maximum of 8 in order to account for sites with higher scores possibly leading to target-directed miRNA degradation (TDMD).

The repression by miRNA $g$ can then be understood as the ratio between its occupancy and a background occupancy term, in which the dissociation constants are set to that of nonspecifically bound sites (i.e. $K_d = 1.0$). More specifically, McGeary et al. (2019) model the repression as:

$$N_{m,g,\mathrm{background}} = \sum_{i=1}^{p} \left( \frac{a_g}{a_g + c_{\mathrm{ORF}}} \right) + \sum_{j=1}^{q} \left( \frac{a_g}{a_g + 1} \right)$$

where $b$ can be interpreted as the additional repression caused by a single bound AGO. Because UTR and ORF lengths have been reported to influence the efficacy of repression (Agarwal et al., 2015; Hausser et al., 2009), scanMiR allows for an additional term to take these effects into account:

$$\text{repression}_{\text{adj}} = \text{repression} \cdot (1 + f \cdot \text{UTR.length} + h \cdot \text{ORF.length})$$

*UTR.length* and *ORF.length* are linearly normalized so that 0 and 1 are respectively the 5% and 95% quantiles of the distribution of lengths (see Garcia et al. 2011; Agarwal et al. 2015). This adjustment overall leads to slightly improved correlations with observed repression (Suppl. Fig. 7B-C). Given that the effect is small except for some extreme transcripts, these parameters are however set to 0 by default.

While $b$, $c$, $p3$, $f$ and $h$ are considered global parameters (i.e. the same for different miRNAs and transcripts and also across experimental contexts), $a$ is expected to be different for each miRNA in a given experimental condition. Values for parameters $b$ (= 1.77) and $c$ (= -1.71) were therefore obtained from the Biochemical Plus model of McGeary et al. (2019), which was optimized with the experimentally determined RBNS data. Parameters $p3$, $f$ and $h$ were globally fitted to maximize the correlation with miRNA transfection experiments from McGeary et al. (2019).

As shown by McGeary et al. (2019), the performance of the biochemical model is robust to changes in parameter $a$ of several orders of magnitude (see also Suppl. Fig. 3A-B). Furthermore, repression predictions obtained with the globally optimized $a$ value still outperform TargetScan substantially (see Fig. 3 & Suppl. Fig. 2). Therefore, scanMiR provides reasonable default parameters, although users can easily provide their own parameter values.

### Comparison of predicted and observed repression

For HeLa and HEK cell lines, we used the transcriptome reconstructions and quantifications provided by the authors in the GEO series (respectively in series GSE140217 and GSE140218). Given the absence of controls, both expression and predicted repression were normalized as described in McGeary et al. (2019). TargetScan7 values were obtained by running the TargetScan7 Python scripts of Kathy Lin (https://github.com/kslin/targetscan) with default parameters and conservation files supplied in the Git repository. The 100way Multi-Species Alignment (MAF) file of the custom reconstructed HEK transcripts of McGeary et al. (2019) was obtained by downloading alignments from the UCSC Genome Browser with mafFetch and then further processing these with "Stitch MAF blocks" from Galaxy (Blankenberg et al., 2011) and R. Common species identifiers were obtained from the NCBI Taxonomy Browser. miRNA seed and family information was downloaded from the TargetScan7 homepage (http://www.targetscan.org/vert_72/) and further processed in R to conform to the example input files. TargetScan8 occupancy scores were downloaded from the TargetScan8 homepage (http://www.targetscan.org/vert_80/).

To assess the scanMiR performance in another species, we downloaded two datasets of miRNA knockout studies performed in mice (Amin et al., 2015; Eichhorn et al., 2014). Reads were mapped to the GRCm38 genome and subsequently counted with Salmon 1.3.0 (Patro et al., 2017). logFC-values were obtained from edgeR (v. 3.32) (Robinson et al., 2010), by filtering for expressed transcripts using the filterByExpr-function and normalizing with a weighted trimmed mean of the logarithmic expression ratios of individual samples (TMM). For the miR-122 knockout dataset from Eichhorn et al. (2014), a common negative binomial dispersion was estimated since the authors performed RNA-sequencing with only one replicate. In order to correlate logFCs of these two datasets with scanMiR repression predictions, we considered only transcripts that constitute 90% of the expressed transcripts of one gene in the specific setting, that are expressed higher than 10 TPM (Amin et al., 2015) or 0.5 TPM (Eichhorn et al., 2014), respectively, that are reported as representative transcripts in TargetScan, and that are supported with at least five 3p-seq-tags (http://www.targetscan.org/mmu_80/mmu_80_data_download/Gene_info.txt.zip). Mouse TargetScan8 scores were downloaded from http://www.targetscan.org/mmu_80/ and are based on the custom TargetScan mouse 3'UTR annotations.

#### Other external datasets used

miRNA expression changes upon TDMD knockout in induced mouse neurons at day 10 of differentiation were downloaded from the supplementary data (Data S2) from Shi et al. (2020). Expression changes with an adjusted p-value smaller than 10-5 were considered significant (concurrent with the 17 significantly changing miRNAs in Fig. 4B of Shi et al. 2020). Corresponding transcript expression levels were obtained from the study of Whipple et al. (2020). Sequenced reads were mapped to GRCm38 and afterwards counted using Salmon (Patro et al., 2017). Putative TDMD sites are shown on transcripts with expression levels of at least 10 TPM in more than one wild-type sample of neurons at day 10 of differentiation (Fig. 5A, Supplementary Table 1)

For the circular RNA scans, we used full circular RNA reconstructions from Zhang et al. (2021). A gtf file of the coordinates was kindly provided by the authors, containing also back-splice junction (BSJ) counts. We extracted the corresponding spliced sequences, appending the first 11 nucleotides at the end to enable the identification of sites spanning the back-splice junction. miRNA expression levels in the brain (Fig. 5B, Fig. 6B) were calculated from the supplementary information of Chiang et al. (2010).

#### References

Agarwal, V., Bell, G. W., Nam, J. W., & Bartel, D. P. (2015). Predicting effective microRNA target sites in mammalian mRNAs. eLife, 4, 1–38. https://doi.org/10.7554/eLife.05005

Amin, N. D., Bai, G., Klug, J. R., Bonanomi, D., Pankratz, M. T., Gifford, W. D., Hinckley, C. A., Sternfeld, M. J., Driscoll, S. P., Dominguez, B., Lee, K. F., Jin, X., & Pfaff, S. L. (2015). Loss of motoneuron-specific microRNA-218 causes systemic neuromuscular failure. Science, 350(6267), 1525–1529. https://doi.org/10.1126/science.aad2509

Chiang, H. R., Schoenfeld, L. W., Ruby, J. G., Auyeung, V. C., Spies, N., Baek, D., Johnston, W. K., Russ, C., Luo, S., Babiarz, J. E., Blelloch, R., Schroth, G. P., Nusbaum, C., & Bartel, D. P. (2010). Mammalian microRNAs: Experimental evaluation of novel and previously annotated genes. Genes & Development, 24(10), 992–1009. https://doi.org/10.1101/gad.1884710

Eichhorn, S. W., Guo, H., McGeary, S. E., Rodriguez-Mias, R. A., Shin, C., Baek, D., Hsu, S., Ghoshal, K., Villén, J., & Bartel, D. P. (2014). MRNA Destabilization Is the Dominant Effect of Mammalian MicroRNAs by the Time Substantial Repression Ensues. Molecular Cell, 56(1), 104–115. https://doi.org/10.1016/j.molcel.2014.08.028

Garcia, D. M., Baek, D., Shin, C., Bell, G. W., Grimson, A., & Bartel, D. P. (2011). Weak seed-pairing stability and high target-site abundance decrease the proficiency of lsy-6 and other microRNAs. Nature Structural & Molecular Biology, 18(10), 1139–1146. https://doi.org/10.1038/nsmb.2115

Grimson, A., Farh, K. K. H., Johnston, W. K., Garrett-Engele, P., Lim, L. P., & Bartel, D. P. (2007). MicroRNA Targeting Specificity in Mammals: Determinants beyond Seed Pairing. Molecular Cell, 27(1), 91–105. https://doi.org/10.1016/j.molcel.2007.06.017

Hausser, J., Landthaler, M., Jaskiewicz, L., Gaidatzis, D., & Zavolan, M. (2009). Relative contribution of sequence and structure features to the mRNA binding of Argonaute/EIF2C–miRNA complexes and the degradation of miRNA targets. Genome Research, 19(11), 2009–2020. https://doi.org/10.1101/gr.091181.109

McGeary, S. E., Lin, K. S., Shi, C. Y., Pham, T. M., Bisaria, N., Kelley, G. M., & Bartel, D. P. (2019). The biochemical basis of microRNA targeting efficacy. Science, 366(6472). https://doi.org/10.1126/science.aav1741

Blankenberg, D., Taylor, J., Nekrutenko, A., & The Galaxy Team. (2011). Making whole genome multiple alignments usable for biologists. Bioinformatics, 27(17), 2426–2428. https://doi.org/10.1093/bioinformatics/btr398

Patro, R., Duggal, G., Love, M. I., Irizarry, R. A., & Kingsford, C. (2017). Salmon provides fast and bias-aware quantification of transcript expression. Nature Methods, 14(4), 417–419. https://doi.org/10.1038/nmeth.4197

Robinson, M. D., McCarthy, D. J., & Smyth, G. K. (2010). edgeR: A Bioconductor package for differential expression analysis of digital gene expression data. Bioinformatics, 26(1), 139–140. https://doi.org/10.1093/bioinformatics/btp616

Shi, C. Y., Kingston, E., Kleaveland, B., Lin, D. H., Stubna, M. W., & Bartel, D. P. (2020). The ZSWIM8 ubiquitin ligase mediates target-directed microRNA degradation. Science, 21(1), 1–9. https://doi.org/10.1126/science.abc9359

Whipple, A. J., Breton-provencher, V., Jacobs, H. N., Chitta, U. K., Sur, M., Sharp, P. A., Whipple, A. J., Breton-provencher, V., Jacobs, H. N., Chitta, U. K., & Sur, M. (2020). Imprinted Maternally Expressed microRNAs Antagonize Paternally Driven Gene Programs in Article Imprinted Maternally Expressed microRNAs Antagonize Paternally Driven Gene Programs in Neurons. Molecular Cell, 1–11. https://doi.org/10.1016/j.molcel.2020.01.020

Zhang, J., Hou, L., Zuo, Z., Ji, P., Zhang, X., Xue, Y., & Zhao, F. (2021). Comprehensive profiling of circular RNAs with nanopore sequencing and CIRI-long. Nature Biotechnology, 39(7), 836–845. https://doi.org/10.1038/s41587-021-00842-6
